# Targeting Heparan Sulfate 3-O-Sulfation Enhances Acetylcholine Receptor Clustering and Stabilizes Neuromuscular Junctions in Duchenne Muscular Dystrophy

**DOI:** 10.64898/2026.11.18.767572

**Authors:** Félix Dumont, Ankita Chandra, Alexandre Dobbertin, Frédéric Auradé, Sidwell Bobe Rigade, Saurabh Anand, Alessio Rotini, Marianne Gervais-Taurel, Fanny Coulpier, Mégane Lemaitre, Toin H. van Kuppevelt, Laurent Tiret, Raghavendra Kikkeri, Frédéric Relaix, Olivier Stettler

## Abstract

Duchenne muscular dystrophy (DMD) is a severe neuromuscular disease caused by dystrophin deficiency, leading to progressive muscle degeneration and neuromuscular junction (NMJ) instability. Using rat models of DMD, we identified a dysregulation in heparan sulfate (HS) biosynthesis at the NMJ, characterized by reduced expression of *HsCst2* and *Hs3st5* genes that encode sulfotransferases required for the production of rare 3-O-sulfated heparan sulfate (3S-HS) forms. These genes act synergistically to enhance agrin signaling: their co-expression in myotubes potentiates acetylcholine receptor clustering, and HSPG mimics bearing tetrasulfated 3-O-sulfated motifs reproduce this etfect and promote Agrin-LRP4 interaction. In DMD rats, muscle co-transduction of *HsCst2* and *Hs3st5* prevents NMJ fragmentation and improves muscle strength. These findings identify 3S-HSs as novel regulators of NMJ stability and support HS modulation as a complementary therapeutic strategy in DMD.

## Introduction

Duchenne muscular dystrophy (DMD) is a common and severe muscular dystrophy. X-linked mutations in the *DMD* gene abolish functional dystrophin, which stabilizes the sarcolemma of the muscle fiber. DMD atfects approximately one in 5,000 boys at birth^1^ and causes progressive muscle degeneration and premature death. The most promising therapies currently in clinical trial restore dystrophin function through truncated forms, the micro-dystrophins (for recent reviews see refs^2,3^), or through split-reassembled longer forms of dystrophin^4,5^. Stabilizing NMJ integrity represents a potential important complementary therapeutic approach in DMD because i) NMJ fragmentation impairs maintenance of the postsynaptic architecture and contributes to altered neurotransmission and muscle fatigue, in animal models and in patients^6–9^ and ii) dystrophin replacement may require levels of expression that remain insutficient to fully reverse this fragmentation^10,11^. Loss of dystrophin disrupts the dystrophin-associated protein complex (DAPC) and impairs agrin signaling through the agrin-LRP4-MuSK pathway, a key regulator of nicotinic acetylcholine receptor (AChR) clustering and NMJ stability^12,13^. Accordingly, enhancing agrin signaling through LRP4, mini-agrin, MuSK or downstream etfectors improves NMJ organization and muscle function in the mdx mouse, a model of DMD^14–17^. Agrin signaling is therefore a therapeutic target and raises the question of which extracellular mechanisms regulate it at the NMJ.

Heparan sulfates (HS) emerged as promising candidates, because HS sulfation modulates the interaction of HS with agrin^18,19^. HS are glycosaminoglycans whose diverse sulfation patterns confer selective protein-binding properties^20–23^. As components of heparan sulfate proteoglycans (HSPGs), HS regulate the distribution and signaling of numerous extracellular ligands, and contribute to synaptic organization and function^24,25^. Agrin is itself a large HSPG, and can interact with HS chains carried by other HSPGs at the NMJ^18,19^. The HS structures and the enzymatic pathways involved in NMJ organization nonetheless remain poorly understood. Among HS modifications, 3-O-sulfation generates 3S-HS species that are rare, yet responsible for protein interactions critical to HS function^20,26–29^. We previously showed that 3S-HS-dependent interactions regulate synaptic assembly and activity in vertebrate neurons^30^, a finding since supported independently^31^. Notably, 3-Osulfation is required for NMJ stabilization in the invertebrate *C. elegans*^32^. We therefore assumed that specific 3S-HS and the sulfotransferases that generate them could constitute an additional level of regulation for synaptic organizers such as agrin, in vertebrates. Here we characterized their expression at the normal and the pathological NMJ. Using newly established DMD rat models that recapitulate key features of the human disease^33,34^, and combining transcriptomic, glycobiological, *in vitro* and *in vivo* approaches, we identified *HsCst2* and *Hs3st5* as NMJ-enriched HS sulfotransferase genes whose expression is markedly reduced in DMD. The two encoded enzymes act sequentially to generate 3S-HS, which we detected at the NMJ. *HsCst2* and *Hs3st5* cooperatively potentiate agrin-dependent AChR clustering and prevent NMJ fragmentation in DMD rats *in vivo.* These findings identify 3S-HS as a previously unrecognized regulatory axis of agrin signaling and NMJ stability and suggest that modulating this pathway may preserve the NMJ in DMD.

## Results

Deletion and duplication are the two major classes of mutation underlying human DMD. We therefore used the Del 52 rat model, carrying an exon 52 deletion^33^, and the Dup 10-17 model^34^, carrying a duplication of exons 10-17 in the *Dmd* gene. Both models lose full-length muscle dystrophin (Dp427m) entirely, develop severe striated muscle dystrophy, and die prematurely at approximately 11 months. Using both models supports the identification of mutation-agnostic mechanisms.

We first characterized NMJ morphology in DMD rats using 3D transparency analysis of the tibialis anterior (TA) muscle (Fig. 1). The number of main motor axonal trunks, labeled by neurofilament (NF), was unchanged in both DMD models compared with wild-type (WT) animals (insets in Fig. 1a; Fig. 1b). In contrast, presynaptic morphology (NF/SV2, Fig. 1a, c) and postsynaptic morphology (*α*-bungarotoxin against AChR, Fig. 1a, d) showed considerable fragmentation between 2 and 7 months in both models, consistent with reports in the mdx mouse^14^. Three-dimensional analysis showed the fragmented pre- and postsynaptic compartments to be perfectly superimposed (Fig. 1a), with an enlargement of the motor endplate between 2 and 7 months (Fig. 1a, e). The synaptic volume, defined as the volume of overlap between the two compartments, remained unchanged in all conditions (Fig. 1a, f). Fragmented nerve-muscle contacts therefore retained their overall volume in DMD.

**Fig. 1.**
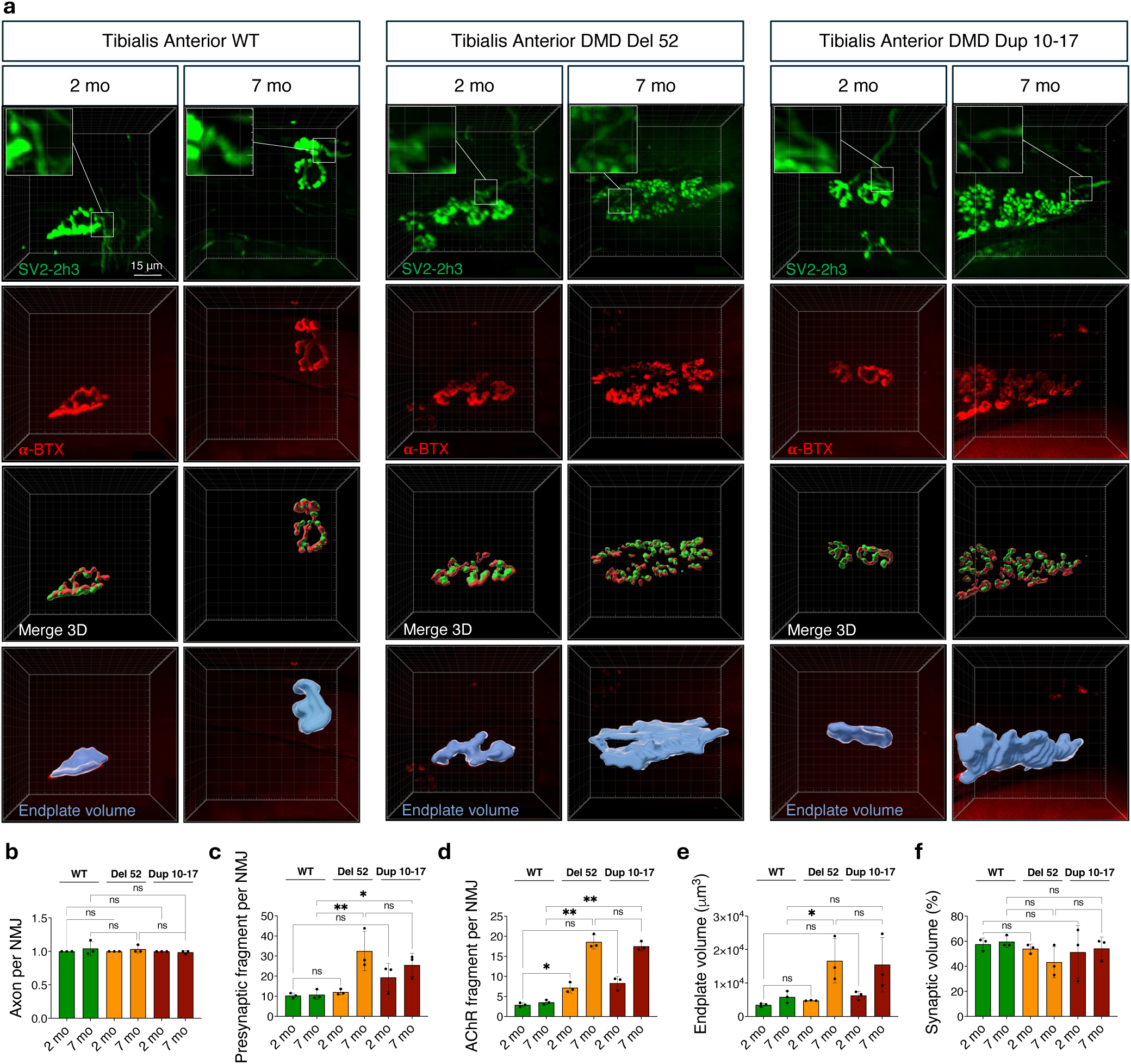
Morphological analysis of NMJ in WT and DMD rat models. **a**, Representative images of NMJ in wild-type (WT) and DMD rats (Del 52 and Dup10-17) aged 2 and 7 months. Axons and presynaptic terminals were co-labeled using an antineurofilament antibody (2h3, axons) and an anti-synaptic vesicle 2 antibody (SV2, presynapse) simultaneously (in green), while endplate was labeled with α-bungarotoxin recognizing AChRs (α-BTX, in red). Morphometric analyses of NMJs were performed based on 3D reconstructions of the NMJs. **b**, Quantification of innervation, assessed by the number of axons in contact with each motor endplate. **c**, Quantification of presynaptic integrity, assessed by the number of SV2 immunoreactive cluster fragments per motor endplate. **d**, Quantification of postsynaptic integrity, assessed by the number of AChR cluster fragments per motor endplate. **e**, Quantification of NMJ spatial dispersion, assessed by motor endplate volume (in blue). **f**, Quantification synaptic volume characterized as pre- and postsynaptic colocalization volume. Data are derived from n = 3 animals per condition, with 30 NMJs analyzed per animal. Mean ± s.e.m. ; ns, not significant; *P < 0.05; **P < 0.01. Scale Bar, 15 μm. Bar applies for all panels.

To examine the molecular basis of the DMD NMJ, we performed comparative single-nucleus RNA sequencing (snRNA-seq) of muscle nuclei from 11-month-old Del 52 and control rats. To enrich the samples in subsynaptic myonuclei, we prepared nuclear suspensions from the medial regions of the diaphragm, an area of high NMJ density^35^ (Fig. 2). UMAP plots were generated from 6,771 WT nuclei and 4,374 DMD nuclei, with 39,000 and 58,000 mean reads per nucleus and approximately 2,500 and 352 genes per nucleus, respectively. Integration of the rat datasets distinguished fifteen nucleus types by expression profile (Fig. 2a), including body myonuclei, NMJ myonuclei, Schwann cells, satellite cells, fibroblast and fibro-adipogenic progenitors (FAPs), immune cells, smooth muscle cells, endothelial cells, and adipocytes. DMD clusters resembled those reported in mice carrying a DMD deletion^36^, with altered proportions of myogenic lineage nuclei, NMJ nuclei excepted, and generalized transcriptome alterations (Fig. S1a, b).

**Fig. 2.**
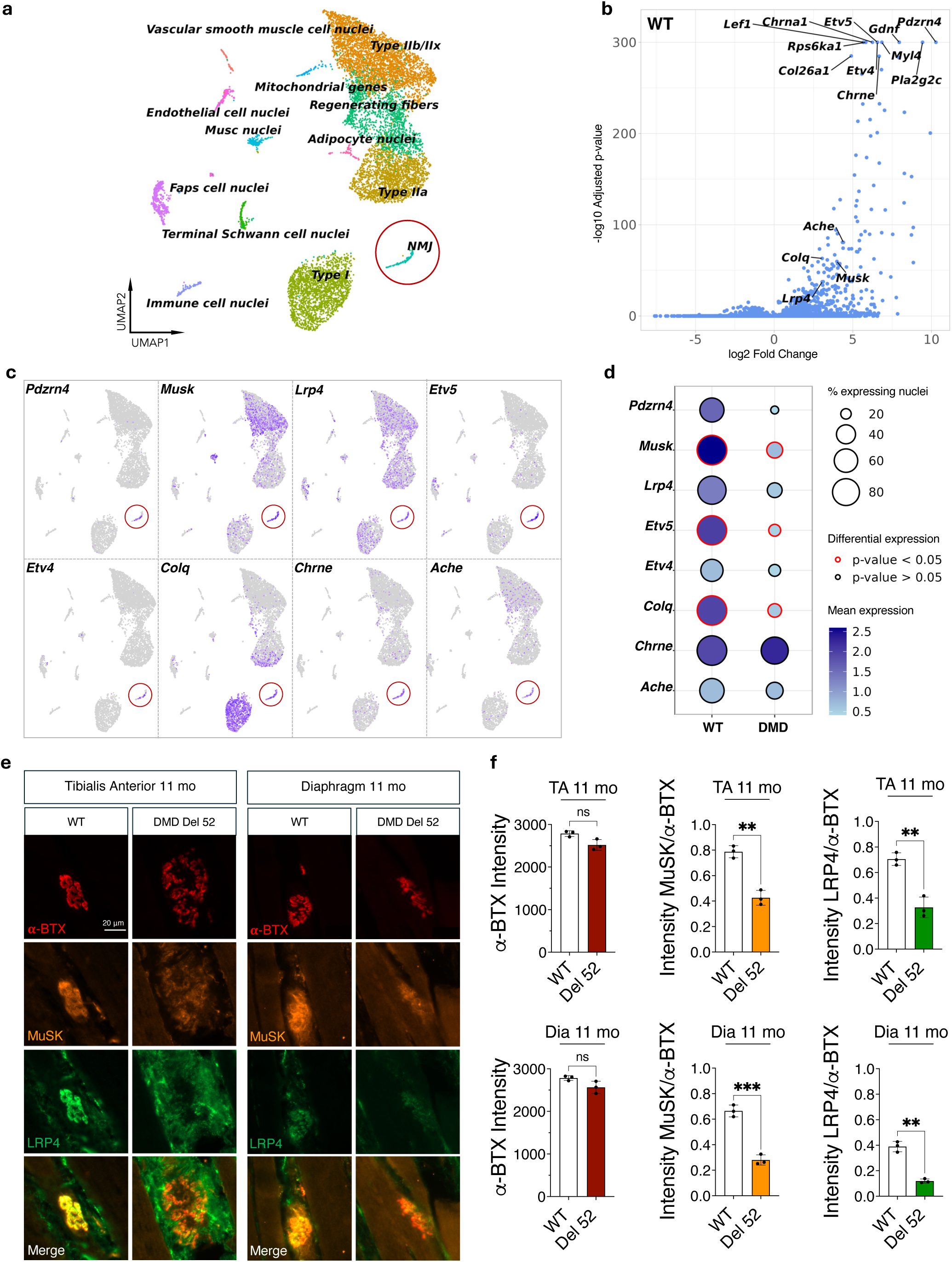
Comparative transcriptomic analysis of key NMJ genes in WT and DMD skeletal muscle. **a**, UMAP representation of myonuclei clusters derived from snRNA-seq analyses of skeletal muscle. The NMJ myonuclei cluster is circled in red. **b**, Volcano plot showing NMJ genes (e.g., *Pdzrn4*, *Musk*, *Lrp4*, *Etv4/5*, *ColQ*, *Chrne*, *AChE*) enriched in the WT NMJ cluster compared to other clusters. **c**, Feature plots illustrating the enrichment of NMJ marker genes within the NMJ cluster. **d**, Dot plot comparing the expression of NMJ marker genes between WT and DMD NMJ clusters. **e**, Representative fluorescent images of NMJs from 11-month-old WT and DMD rat *tibialis anterior* and *diaphragm* muscles co-labeled with α-bungarotoxin (α-BTX, red), anti-MuSK antibody (in orange), and anti-LRP4 antibody (in green). Scale bar, 20 μm. Bar applies for all panels. **f**, Quantification of α-bungarotoxin labeling-normalized immunofluorescence of MuSK and Lrp4 in NMJ of the tibialis anterior and diaphragm. Data are from n = 3 animals per condition, with 70 NMJs analyzed per animal. Mean ± s.e.m.; statistical comparisons were performed using an unpaired two-tailed t-test: ns, not significant; *P < 0.05; **P < 0.01; ***P < 0.001; ****P < 0.0001.

We first validated the NMJ cluster, initially defined in WT animals with a limited set of standard markers (*Colq*, *Ache, Chrne, Lrp4).* This cluster captured those canonical genes and was further enriched in NMJ-related genes, including markers previously detected by snRNA-seq (Musk, *Chrna1, Chrnd, Etv4, Etv5, Pdzrn4)*^37,38^ and genes more recently associated with the NMJ (e.g. Rps6ka1)^37,39,40^ (Supplementary File 1; Fig. 2b, c). Within the NMJ gene group, DMD condition showed downregulation of genes involved in NMJ maintenance (*Pdzrn4*), agrin signaling (*Lrp4*, *Musk),* acetylcholine degradation (*Ache*, *Colq),* and synaptic transcription (*Etv4*, *Etv5*) (Fig. 2d). *Chrne*, which encodes the mature epsilon subunit of the AChR, did not change, suggesting that the capacity to synthesize cholinergic receptors remained intact (Fig. 2d). Immunofluorescence against LRP4 and MuSK revealed reduced expression of both proteins in the TA and diaphragm of 11-month-old Del 52 rats (Fig. 2e, f), consistent with the snRNA-seq data and validating the dataset.

The implication of HS in NMJ stability in the invertebrate C. *Elegans*^32^ prompted us to search for HS biosynthesis genes enriched at the WT NMJ and downregulated in DMD. We identified *HsCst2* and *Hs3st5* (Fig. 3a-d). These genes respectively encode HS 6-O-sulfotransferase 2 (HS6ST2) and HS 3-O-sulfotransferase 5 (HS3ST5), which are required, among other members of their respective families^20^, for the terminal 3-O-sulfation of HS (Fig. S2) and for the production of 3S-HS with specific protein-binding capabilities^28,41^. Analysis of the MyoAtlas dataset^39^ further revealed that mouse NMJs are enriched in *HsCst2* and in the *Hs3st* family members *Hs3st3a1* and *Hs3st3b1* (Fig. S3). *Hs3st5* was expressed at mouse NMJs without reaching the enrichment seen in rats (https://research.cchmc.org/myoatlas/). These findings suggested a conserved sulfotransferase signature in rodent NMJ. The reduced expression of *HsCst2* and *Hs3st5* in DMD rats (Fig. 3b-d) suggested a reduced capacity to synthesize 3S-HS. This reduction was compounded by the parallel downregulation of other genes required for HS assembly, which are not enriched at the NMJ (Fig. S2, Fig. 3d). These genes act at every step of chain construction, from initiation (*Xylt1-2*) and polymerization (*Ext1, Ext2*) to N-sulfation (*Ndst1-4*), epimerization (*Glce*), and 2-O-sulfation (*Hs2st1*). *Papss2* was the only HS biosynthetic gene trending upward (Fig. 3b, d). Papss2 encodes 3′-phosphoadenosine 5′-phosphosulfate synthetase 2, which produces the sulfate donor PAPS, and we interpreted this trend as a possible compensatory response to the loss of HS-dependent signaling. Over-representation analysis against the Gene Ontology database gave further support to a central role for HS at the NMJ, since glycosaminoglycan binding and heparin binding ranked among the key molecular functions of the WT NMJ gene cluster and were absent from the DMD cluster (Fig. S4).

**Fig. 3.**
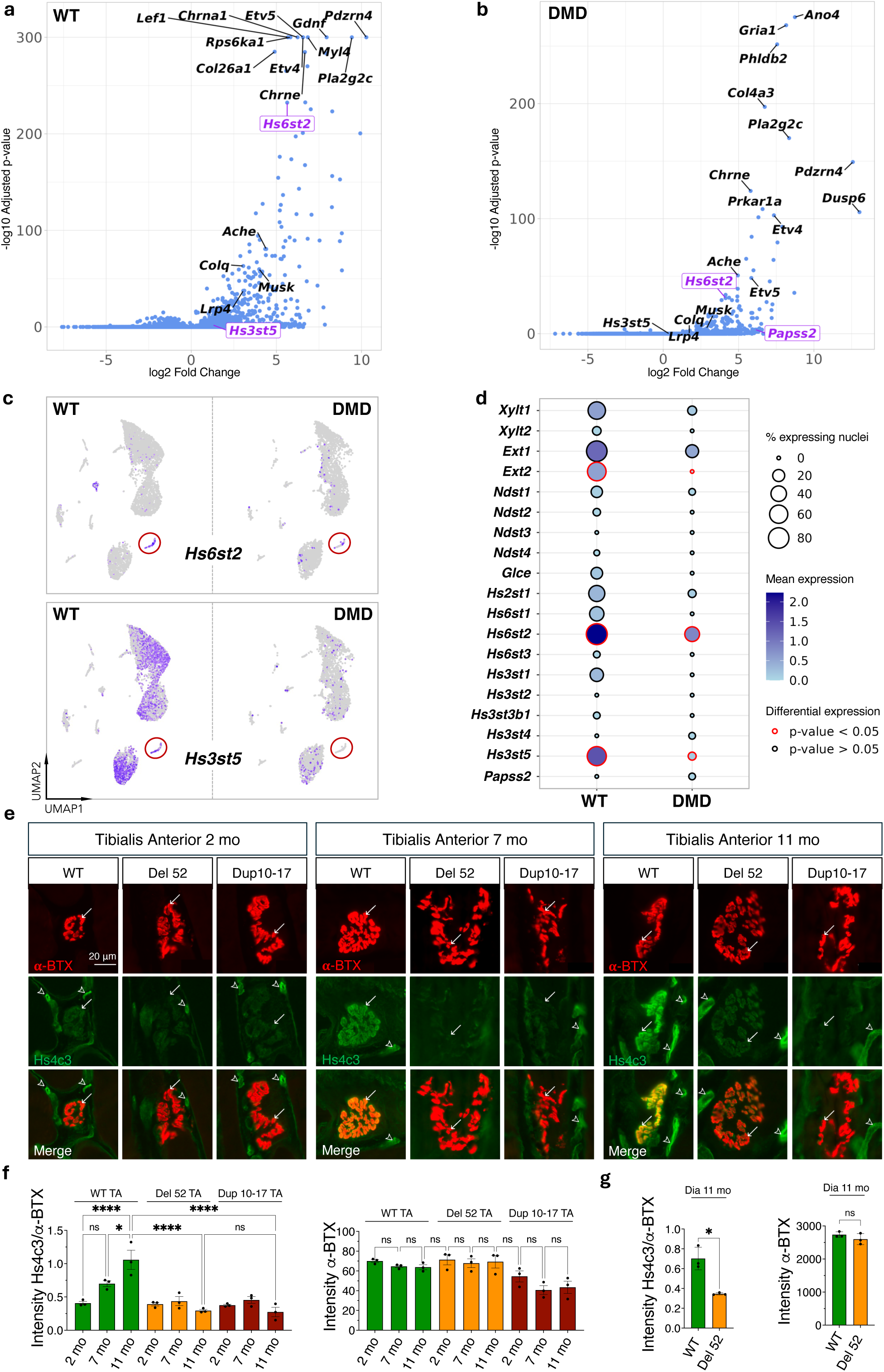
Dysregulation of heparan sulfate biosynthesis pathway at the DMD rat NMJ. **a**, Volcano plot of muscle WT genes differentially expressed between NMJ and non-NMJ myonuclei, highlighting in purple *Hs6st2* and *Hs3st5* among NMJ-enriched genes. **b**, Volcano plot of genes enriched in NMJ myonuclei in DMD. **c**, Feature plots comparing *Hs6st2* and *Hs3st5* expression in WT versus DMD myonuclei. **d**, Dot plot of heparan sulfate biosynthesis genes in WT versus DMD myonuclei. **e**, Representative fluorescent images of NMJ from 2, 7, and 11 months WT and DMD tibialis anterior muscle labeled with α-bungarotoxin (α-BTX, in red) and Hs4c3 antibody recognizing preferentially 3-O-sulfated heparan sulfate (3S-HS) chain (in green). White arrows point to NMJ and arrowheads to blood vessels. Scale bar, 20 μm. Bar applies to all panels. **f**, Quantification of Hs4c3 fluorescence intensity normalized to α-bungarotoxin labelling in *tibialis anterior* across ages and genotypes. **g**, Same quantification than in (f), but for the *diaphragm* at 11 months old. Data are from n = 3 animals per condition, with 70 NMJs per animal; mean ± s.e.m.; statistical comparisons were performed using a Kruskal-Wallis with Dunn’s correction (f) or unpaired two-tailed t-test (g); ns, not significant; * P < 0.05; ****P <0.0001.

To validate this transcriptomic identification, we sought to detect 3S-HS at the NMJ, since these HS are the products of the sequential activity of HS6ST2 and HS3ST5 (Fig. S2). We used the Hs4c3 antibody, which preferentially recognizes 3S-HS^42^. Hs4c3 labeled NMJs in adult rats and mice (Fig. 3e, Fig. S5), and additional labeling coincided with blood capillaries throughout the muscle (Fig. 3e). Hs4c3 has occasionally been reported to recognize HS chains bearing NS2S6S sequences lacking 3-O-sulfate groups^42^, but its reactivity here tracked 3-O-sulfotransferase expression (see below) and therefore the presence of 3S-HS. At the NMJ, most Hs4c3 labeling co-localized with AChR identified by *α*-BTX (Fig. S5). In WT rats, Hs4c3 labeling at the NMJ rose progressively, first between 2 and 7 months and again between 7 and 11 months, indicating increased 3S-HS production as the adult NMJ matures (Fig. 3e, f). In DMD rats, Hs4c3 labeling at 7 and 11 months remained as low as at 2 months (Fig. 3e, f). Defective maintenance of 3S-HS at the NMJ therefore correlated with the downregulation of the mRNAs encoding the enzymes that produce 3S-HS.

Agrin, the major NMJ organizer, binds multisulfated HS sequences on synthetic oligosaccharide microarrays^19^. We therefore asked whether *HsCst2* and *Hs3st5* atfect agrin signaling through 3S-HS, using C2C12 myotubes. Given the low basal expression of both *genes* in C2C12 (Fig. S6), we overexpressed them individually and in combination, by co-transfection, before myoblast fusion into myotubes. Transfected myotubes were then treated with a glycan-free recombinant neuronal agrin (Fig. 4a). Simultaneous overexpression of *HsCst2* and *Hs3st5* significantly enhanced agrin activity, increasing the number of AChR clusters by approximately 70% over basal activity (Fig. 4b, c). Individual overexpression of each gene, although performed at twice the vector concentration used for co-transfection, produced only a non-significant trend (Fig. 4b, c). Co-transfection also significantly increased agrin-induced cluster length, as did *HsCst2* alone at double vector concentration, unlike *Hs3st5* (Fig. 4b, d). Co-expression of *HsCst2* and *Hs3st5* therefore potentiates both cluster number and cluster length, likely through the generation of 3S-HS on as-yet-unidentified C2C12 HSPGs. Hs4c3 immunolabeling of AChR clusters increased significantly upon co-expression and upon *Hs3st5* expression alone (Fig. 4b, e). *HsCst2* alone produced no such increase, consistent with the preferential recognition of 3S-HS by Hs4c3 and indicating that increased 6-O-sulfation on an N-S and 2-O-S background does not sutfice to generate the Hs4c3 epitope.

**Fig. 4.**
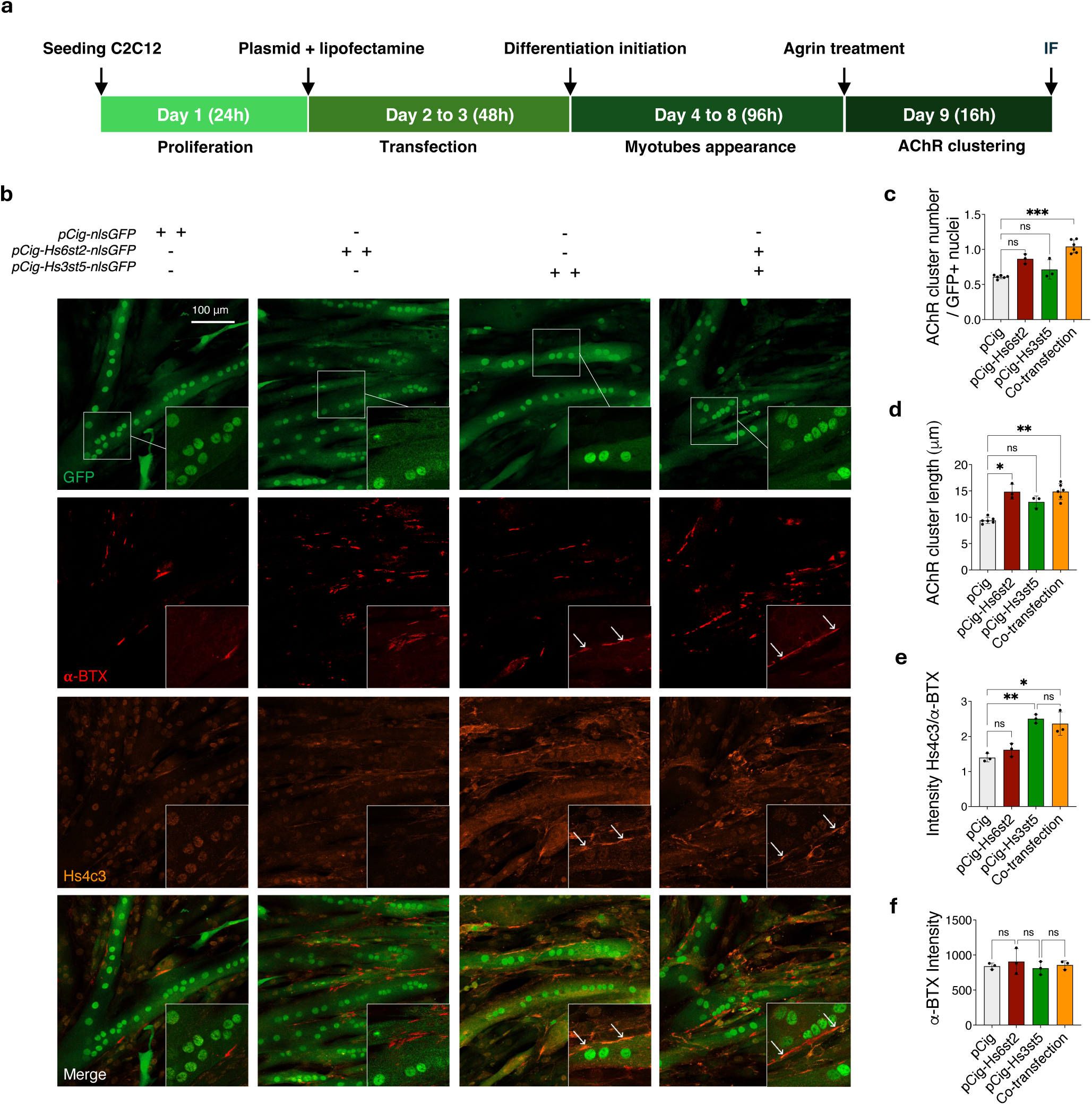
Co-transfection of *Hs6st2* and *Hs3st5* potentiates Agrin-induced AChR clustering in C2C12 myotubes. **a**, Experimental design for the transfection of myoblasts with *Hs6st2* and *Hs3st5* expression vectors and the treatment of the subsequently formed myotubes with Agrin. **b**, Representative images of C2C12 myotubes transfected with pCIG, pCIG*Hs6st2*, pCIG-*Hs3st5*, or co-transfected with both constructs, treated with Agrin and labeled with anti-GFP antibody (in green, transfected nuclei), α-bungarotoxin (α-BTX, in red, AChR), and Hs4c3 antibody (in orange, 3S-HS). Scale bar, 100 μm. Bar applies for all panels. **c**, Quantification of AChR cluster number normalized to GFP⁺ fused myonuclei, (300-600 per independent experiment). **d**, Quantification of AChR cluster length. **e**, Quantification of Hs4c3 antibody immunofluorescence intensity at AChR cluster normalized to α-BTX intensity. **f**, Quantification of α -BTX fluorescence intensity for normalization of HS4C3 labeling. Data are from n = 3-6 independent experiments per condition; Hs4c3 intensity measurements were performed on approximately 100 clusters per independent experiment, mean ± s.e.m. ns, not significant; *P < 0.05; **P < 0.01; ***P < 0.001.

Having shown that combined overexpression of *HsCst2* and *Hs3st5* enhances agrin signaling, we sought the 3S-HS sequences involved. Characterizing HS sequences from cells and tissues remains a challenge in analytical chemistry, in particular for locating the 3-O-sulfate group^20,43^, so we took an indirect approach. We used two naturally occurring oligosaccharide sequences as soluble competitors of the endogenous HSPGs modified by co-transfection, both considered potential products of the sequential activity of HS6ST2 and HS3ST5. The first is an HS3ST3-like product, the gD-like sequence, which forms the binding site for herpesvirus glycoprotein D. The second is an HS3ST1-like product, the AT pentasaccharide, which forms the binding site for antithrombin^20,22,23,28^. HS3ST5 produces both etficiently^20^. We incubated 12-mer oligosaccharides containing AT, gD, or a control sequence lacking 3-O-sulfates (Ctrl)^23^ (Fig. 5a) under basal agrin signaling, without co-transfection, and under enhanced agrin signaling, with co-transfection (Fig. 5b-d). Under basal signaling, none of the three oligosaccharides altered the number of agrin-induced clusters (Fig. 5b, c). In the *HsCst2/Hs3st5* overexpression context, the gD oligosaccharide abolished the supernumerary clusters induced by co-transfection, whereas Ctrl and AT had no etfect (Fig. 5b, c). Free gD oligosaccharides therefore appeared to outcompete for agrin binding the putative gD sequences generated within endogenous HSPGs by co-transfection, while Ctrl and AT were not involved. Cluster length behaved ditferently. Under basal signaling, all three oligosaccharides reduced cluster length relative to agrin alone (Fig. 5b, d), indicating that cluster size under basal agrin activity was driven by the 2SNS6S pattern of Ctrl, without 3-O-sulfates. In transfected cells, the increase in cluster length was partially inhibited by AT and by gD, but not by Ctrl (Fig. 5b, d), a neutralizing etfect of both 3-O-sulfated sequences distinct from the gD-specific control of supernumerary cluster formation. Specific 3S-HS sequences therefore fine-tune distinct modalities of agrin-mediated potentiation of AChR clustering (Fig. 5e).

**Fig. 5.**
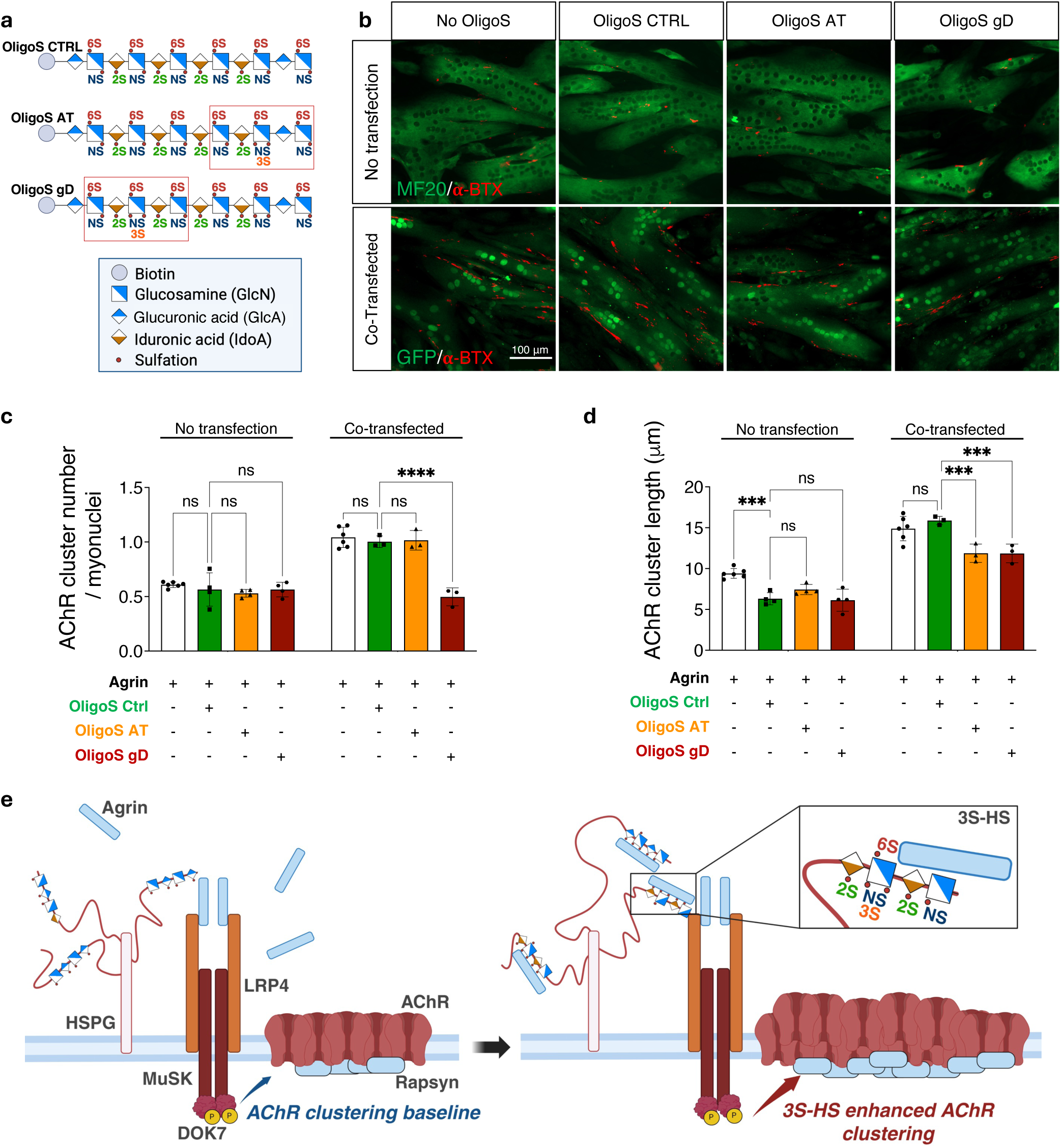
Competition with synthetic 3-O-sulfated oligosaccharides identifies gD- and AT-like sequences as mediators of HS6ST2/HS3ST5-dependent Agrin signaling. **a**, Structures of the synthetic 12-mer heparan sulfate oligosaccharides used in the competition assay: OligoS CTRL (OligoSaccharide without 3-O-sulfate), OligoS AT contains a 3-O-sulfated antithrombinbinding type motif, and OligoS gD, a 3-O-sulfated glycoprotein D-binding type motif. **b**, Representative fluorescent images of C2C12 myotubes labeled with α-bungarotoxin (α-BTX, in red) under basal Agrin signaling (no transfection, counterstain with MF20 antibody, in green) or enhanced Agrin signaling following *Hs6st2*/*Hs3st5* co-transfection (immunolabeled with GFP, in green) in presence or absence of the indicated oligosaccharide competitor. Scalebar, 100 μm. Bar applies for all panels. **c**, Quantification of AChR cluster number normalized to fused myonuclei under basal and enhanced Agrin signaling, per oligosaccharide condition. **d**, Quantification of AChR cluster length under the same conditions. Data from n = 3-6 independent experiments per condition; mean ± s.e.m. ns, not significant; *P < 0.05; ***P < 0.001; **** P < 0.0001. **e**, Scheme of Agrin signaling pathway (left) and increase of AChR clustering by 3S-HS (right). *Created in BioRender. Relaix, F. (2026)* https://BioRender.com/06nexl4.

The competition experiments suggested that the mode of presentation of the oligosaccharide, proteoglycan-linked or free, modulates agrin signaling. To test this directly, we assessed the etfect of HSPG mimics (neoPGs) bearing ditferentially sulfated saccharides (Fig. 6). We screened a previously constructed library of membrane-integrable neoPGs carrying glycosaminoglycans of varied sulfation profile and chain length^44,45^, focusing on neoPGs co-modified at the 6-O and 3-O positions, M14 and M17 (Fig. 6a). neoPG-M14, previously designated neoPG-H11^44^, bears tetrasaccharides (GlcA2S-GlcNS6S3S-GlcA2S) carrying the tetrasulfated 2SNS6S3S sequence of the natural gD sequence (IdoA2S-GlcNS6S3S-IdoA2S), with glucuronic acid in place of iduronic acid. neoPG-M17 (neoPG-H10 in ref^44^) lacks the 2-O-sulfate group on the glucuronic acid. Incubating neoPG-M14 with agrin increased the number of AChR clusters by 40% over agrin alone (Fig. 6b, c), dose-dependently (Fig. 6d). By contrast, neoPG-M15 and neoPG-M17 reduced the etfect of agrin alone (Fig. 6b, c), and neoPG-M18 had no etfect. The inhibition by M17 indicates that the 2-O-sulfation of the uronic acid adjacent to GlcNS6S3S in neoPG-M14 is required for the additional cluster formation. No potentiation was observed with neoPGs lacking 3-O-sulfate groups (M15, M18) or, conversely, carrying 3-O-sulfate but lacking 6-O-sulfate groups (M16). These results highlight the importance of the 6S3S motif and of its symmetric 2-O-sulfated environment in the potentiation of agrin activity. neoPG-M18 carries a higher negative charge than neoPG-M14 yet failed to atfect agrin signaling, which supports a sequence-specific rather than a charge-dependent mechanism.

**Fig. 6.**
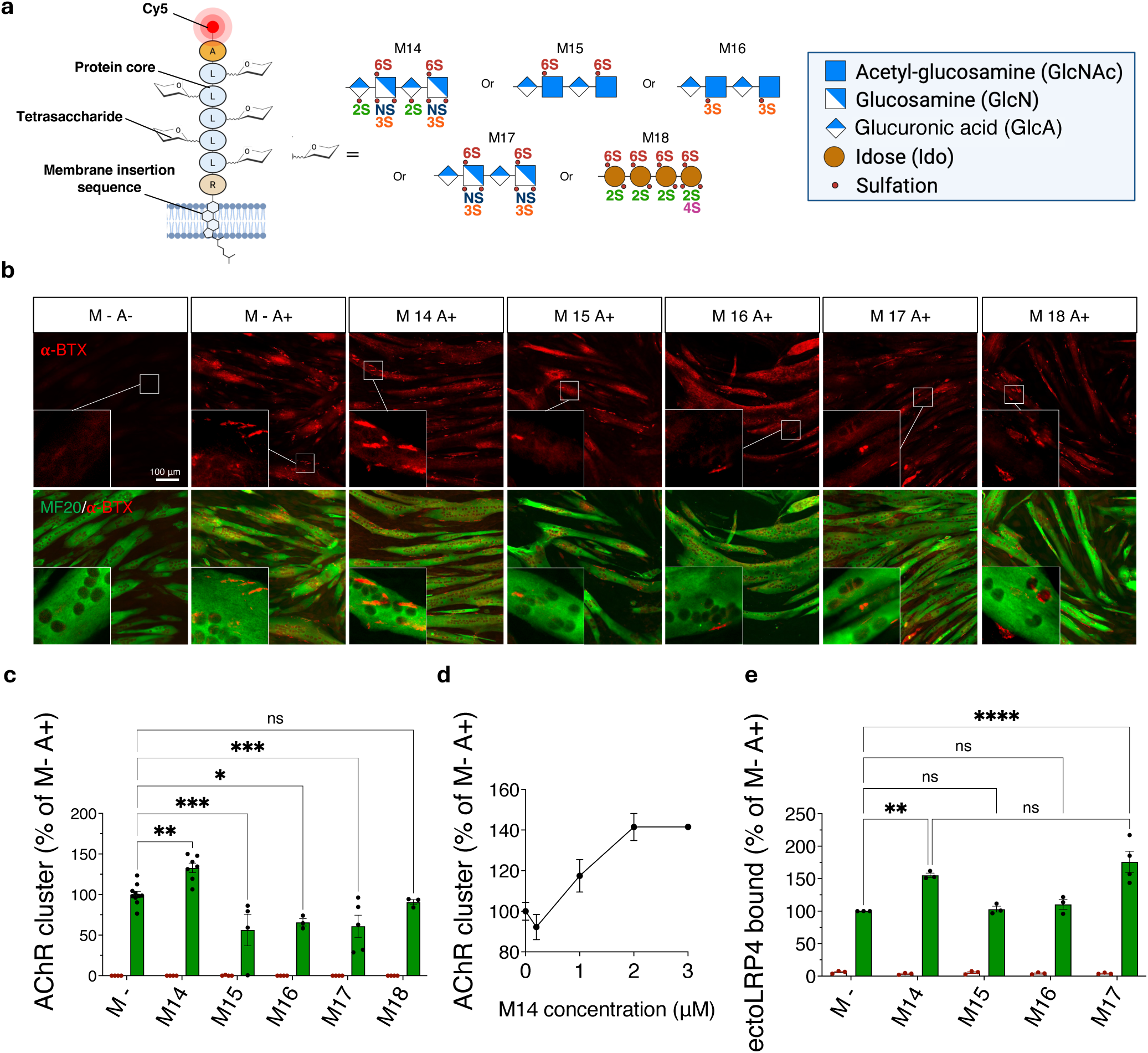
Membrane-insertable neoPGs carrying a gD-type sulfation profile specifically potentiate Agrin-induced AchR clustering. a, Membrane-insertable HSPG mimic structures (neoPGs M14 to M18) constituted of 7 amino acids core bearing five identical tetrasaccharides with varying sulfation profiles as shown in ref 42 , *Created in BioRender. Relaix, F. (2026)* https://BioRender.com/06nexl4. b, Representative images of C2C12 myotubes treated with Agrin alone (M-, A+) or coincubated with each neoPG (M14-M18, A+) and labeled for α-bungarotoxin (α-BTX, in red, AChR) and MF20 (in green). Scale bar, 100 μm. c, Number of AChR clusters under neoPG treatment conditions normalized to number of fused myonuclei, as percentage of the Agrin-only condition. d, Dose-response relationship of Agrin-induced AChR cluster formation with increasing concentrations of neoPG-M14. e, ELISA-like assay illustrating the effect of neoPGs M14-M17 on ectoLRP4 binding to Agrin, normalized to control without NeoPG. Mean ± s.e.m. Statistical comparisons were performed using a two-way ANOVA with Tukey’s multiple comparison test; ns, not significant; *P < 0.05; **P < 0.01; ****P < 0.0001.

Having established that neoPG-M14 recapitulates the etfect of *HsCst2/Hs3st5* co-overexpression on agrin-mediated AChR cluster formation, we investigated the underlying mechanism. Given the pivotal role of LRP4 in initiating agrin signaling^12^, we asked whether neoPG-M14 modulates the agrin-LRP4 interaction. In an ELISA-like assay^46^, neoPG-M14 enhanced agrin binding to LRP4 (Fig. 6e), whereas neoPGs carrying exclusively 6-O-sulfated (M15) or 3-O-sulfated (M16) tetrasaccharides did not. neoPG-M17 enhanced agrin-LRP4 binding despite reducing agrin signaling (Fig. 6e). HS-dependent facilitation of the agrin-LRP4 interaction therefore represents an upstream checkpoint that leads to either potentiation or inhibition of downstream signaling, depending on HS structure.

We next asked whether *HsCst2* and *Hs3st5* cooperative function could prevent NMJ fragmentation *in vivo* in DMD rats. MyoAAV vectors encoding *HsCst2* and *Hs3st5* under the muscle-specific MHCK7 promoter were co-injected into the TA muscles of two-month-old DMD rats, before the apparent onset of NMJ fragmentation (Fig. 7a). Co-transduction produced widespread transgene expression throughout the TA, with a mosaic distribution of nuclei expressing both transgenes, either one, or neither (Fig. S7a, b). Co-and mono-transduced nuclei were treated as equally relevant, since distinct nuclei within one fiber may express ditferent transgenes and collectively supply functional HS6ST2/HS3ST5 pairs in the Golgi where HS are sulfated. Consistent with the *in vitro* findings, co-expression in the DMD TA increased Hs4c3 immunoreactivity relative to DMD-GFP controls (Veh) (Fig. S7c, d). At low magnification, Hs4c3 labeling varied across the cytoplasm of muscle fibers and within basal laminae (Fig. S7c), and higher magnification revealed a significant increase at NMJs (Fig. 7b, c). Three months of *HsCst2/Hs3st5* co-expression prevented NMJ fragmentation, with AChR cluster and presynaptic contact morphology preserved at WT levels (Fig. 7d, e). Co-expression was therefore sutficient to restore AChR cluster organization *in vivo* in DMD rats without additional treatment. To test whether this structural rescue was translated functionally, we measured the *in situ* specific force of the TA after sciatic nerve stimulation immediately before sacrifice. Compared to untreated rats with DMD, co-transduction resulted in a significant improvement in specific muscle strength (i.e., normalized to the mass of the tibialis muscles) (Fig. 7f). Other deleterious parameters of the model^33^ remained unchanged between untreated and treated DMD rats, namely muscle mass (Fig. 7g), the number of regenerating centrally nucleated fibers (Fig. S7a, e), the total number of fibers (Fig. S7f, g), fibrosis (Fig. S7f, i), and fiber size (Fig. S7f, h). These *in vivo* findings corroborate the *in vitro* data, providing proof of concept that 3S-HS enhances AChR clustering. Furthermore, they demonstrate that stimulation of the 3S-HS pathway can improve muscle strength, even in the presence of persistent pathological changes in muscle fibers surrounding the NMJ.

**Fig. 7.**
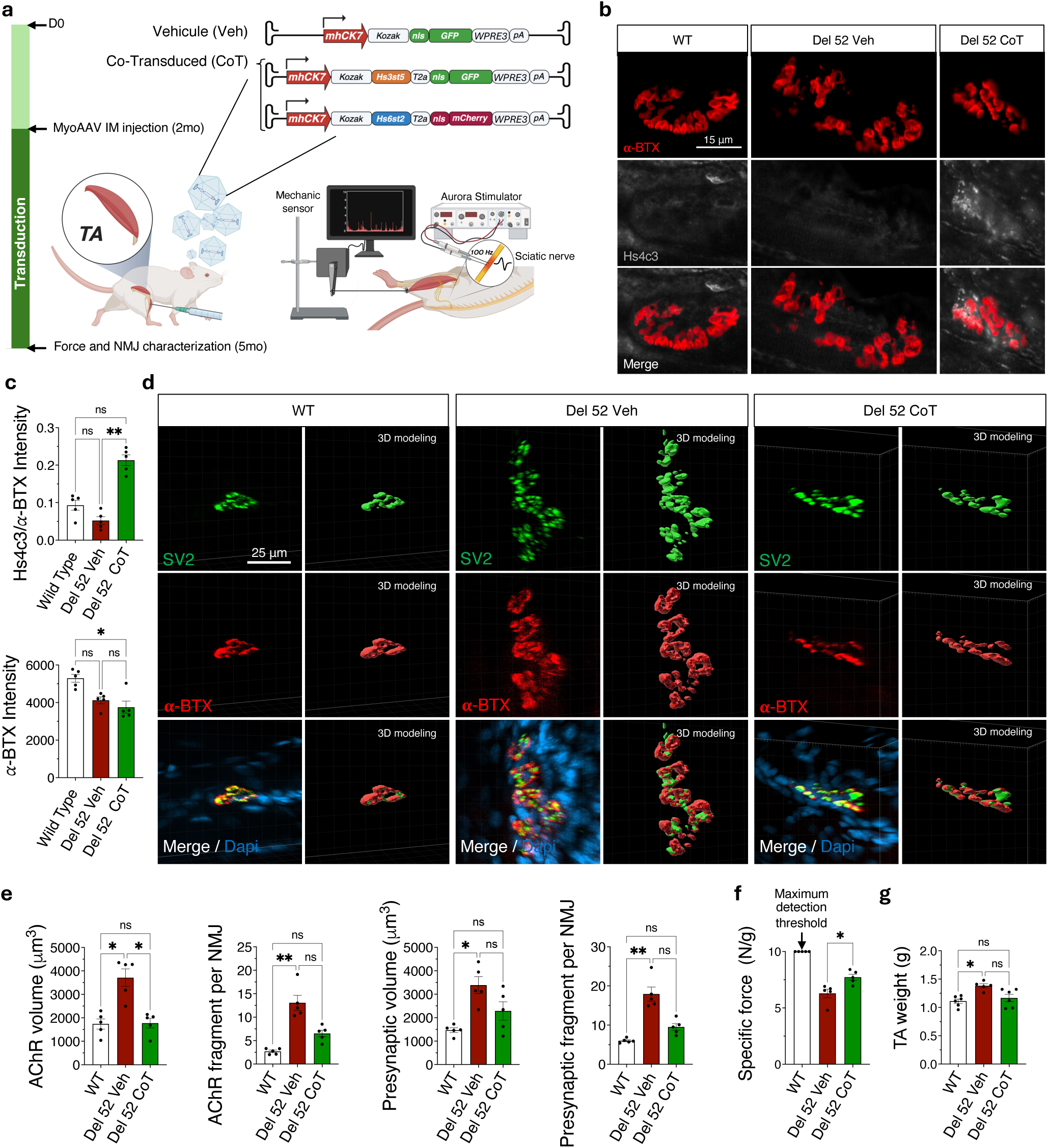
Co-transduction of *Hs6st2* and *Hs3st5* prevents NMJ fragmentation *in vivo*. a, Illustration of the intramuscular administration into the tibialis anterior muscle of Del52 rats of “vehicle” myoAAV vectors or, in combination, *Hs3st5*-T2anlsGFP and *Hs6st2*-T2a-nlsmCherry vectors under the control of the mhCK7 promoter. *Created in BioRender. Relaix, F. (2026)* https://BioRender.com/h4xnjkx. b, Representative NMJ images from WT, Del52 vehicle, and Del52 co-transduced (CoT) TA muscle, labeled for AChR (α-BTX, in red) and 3S-HS (Hs4c3, in grey), with merged images shown below. Scale bar, 25 μm. c, Quantification of Hs4c3 Ab immunofluorescence intensity normalized to α-bungarotoxin at the NMJ. Data from n = 5 animals per group, with 20 NMJs per animal. d, Representative confocal images and 3D reconstructions of NMJs from WT, Del52 vehicle, and Del52 co-transduced rats obtained by tissue clearing, labeled for SV2 (presynaptic, in green), AChR (α-BTX, in red), and merge/DAPI. Scale bar, 25 μm. e, Quantification of endplate (AChR) volume, endplate fragment number, presynaptic (SV2) volume, and presynaptic fragment number, across the three groups. Data from n = 5 animals per group, with 15 NMJs per animal. f, Sciatic nerve stimulation-evoked specific muscle force (N/g) in WT, Del52 vehicle-treated, and Del52 co-transduced rats. g, Tibialis anterior muscle weights used to normalize absolute maximal force. Data are shown as mean ± s.e.m.; each dot presents an individual animal. Statistical comparisons were performed using a Kruskal-Wallis test with Dunn’s multiple comparison correction (c-e;g) or an unpaired two-tailed Student’s t-test (f); *P < 0.05, **P < 0.01; ns, not significant.

## Discussion

Fragmentation of the NMJ, defined by the dispersion of AChR clusters, produces synaptic dysfunction which, combined with muscle fatigue, contributes to muscle weakness in DMD^6,8,9,47^. Here we show that *co-expression* of *HsCst2* and *Hs3st5*, together with their convergent product 3S-HS, potentiates AChR aggregation through an agrin-dependent mechanism. *In vivo,* co-expression preserves NMJ integrity and significantly improves dystrophic muscle strength. These findings identify a synaptic pathway with therapeutic potential in diseases of agrin signaling.

### NMJ co-enrichment of *HsCst2* and *Hs3st5* supports a conserved role for 3S-HS at synapses

Sequential modification of HS by sulfotransferases generates distinct sulfation patterns that create preferential binding sites for specific proteins^20–23^, under tight control by the spatiotemporal expression of individual sulfotransferases^24,48–50^. We identify here the NMJ as a specialized HS microdomain with a distinct sulfotransferase gene signature. *HsCst2* was enriched at the NMJ in rats and in mice. *Hs3st5* was enriched in rats but not in mice, where two other 3-O-sulfotransferase genes, *Hs3st3a1* and *Hs3st3b1,* were enriched instead. Members of both families are therefore enriched in subsynaptic myonuclei, and this signature is conserved across species. HS6ST and HS3ST enzymes act at late steps of the HS sulfation pathway^51^ and collectively generate rare 3S-HS structures with important regulatory functions^20,26,27,29,41^, so their co-enrichment raises the possibility of a synaptic HS sulfation code. Earlier work across species supports this hypothesis. In Drosophila, deletion of the single *HsCst* gene causes structural and functional abnormalities at the NMJ^52^, and in vertebrates *HsCst2* deficiency disrupts postsynaptic organization in the central nervous system^53^. Genetic variants of *Hs3st5* have been associated with autism spectrum disorders and other neuropsychiatric conditions marked by synaptic dysfunction^54^. In *C. elegans,* loss of HS 3-O-sulfation impairs synapse formation in mating-related circuits^55^ and disrupts NMJ organization^32^. We also showed previously that a peptide selectively targeting 3S-HS inhibits synapse formation and alters synaptic function in cultured vertebrate neuronal networks expressing 3-O-sulfotransferases^30^. Our present results extend this conserved role to the vertebrate NMJ. **Co-expression of *HsCst2* and *Hs3st5* generates 3S-HS that potentiates agrin-induced AChR aggregation.**

3S-HS domains interact with a diverse range of ligands and membrane receptors, including FGFR1, neuropilin-1 and agrin^19,20,28,56^, and participate in tissue morphogenesis^23^, coagulation^57^, viral entry^58^, and growth factor signaling^23^. We identify here a role for 3S-HS in potentiating agrin-dependent postsynaptic ditferentiation at the NMJ, and our findings suggest that 3S-HS acts by facilitating the interaction between agrin and LRP4, a key component of the LRP4-MuSK complex that drives AChR clustering. Three independent lines of evidence support a specific role for 3S-HS. First, agrin potentiation is markedly enhanced when 6-O- and 3-O-sulfation are increased simultaneously in myofibers, compared with either pathway alone. Second, only competitive HS oligosaccharides containing 3-O-sulfated motifs abolish the potentiation induced by *HsCst2/Hs3st5* co-overexpression. Third, HSPG mimics containing 3S-HS recapitulate this potentiation. Together these findings link 3S-HS enrichment directly to the potentiation of agrin-induced AChR organization.

Our *in vitro* experiments further reveal a functional distinction between the HS required for basal agrin signaling and the HS responsible for potentiating it. Consistent with Huang and collaborators^19^, we confirm that moderately sulfated HS sutfices to support basal agrin signaling. We identify in addition a potentiating activity of highly sulfated 3S-HS. Two ditferences in experimental design may explain why that potentiating activity was not captured by Huang et al.^19^. First, the low agrin concentration used here may have favored its detection by limiting saturation of AChR clustering. The minimal agrin concentration reported to induce AChR clustering in C2C12 myotubes is 3 pM^59^. We used 30 pM, whereas Huang et al.^19^ used 5 nM in the clustering assay and 50 nM in the HS microarray binding assay, a concentration previously shown to induce AChR clustering in a C2C12 variant deficient in proteoglycan synthesis^59^. Such higher concentrations reduce the dynamic range available for detecting an additional potentiating etfect of HS. Second, Huang et al.^19^ used synthetic polymers displaying approximately 200 copies of a single 2S6S disaccharide, in order to mimic aspects of native HS through multivalent glycan presentation^60^. Our approach instead preserves the endogenous HS environment and selectively increases 3S-HS through *HsCst2/Hs3st5* co-expression. We validated this etfect with a neo-proteoglycan that recapitulates key structural features of endogenous 3-O-sulfated HSPGs, unlike a single-motif glycopolymer. Our competition experiments accordingly assess native, membrane- and matrix-associated HS rather than an artificial polymer presenting one motif at high valency. HS activity at the NMJ therefore depends not only on individual sulfated motifs, but on their combinatorial pattern, sequence context, and native macromolecular organization, with highly sulfated 3S-HS acting as a potentiating component rather than a requirement for basal signaling.

The etfect of 3S-HS on agrin signaling was not uniform. Closely related 3S-HS sequences, the gD- and AT-like motifs, ditferentially regulate distinct features of AChR cluster organization, which suggests that individual 3-O-sulfated motifs confer distinct protein-interaction properties. Cluster number is preferentially regulated by a gD-type motif, whereas cluster size is regulated by both gD- and AT-type sequences. This distinction is unlikely to be incidental, since HS3ST5 is the only HS 3-O-sulfotransferase capable of generating both motifs^20^. The vertebrate family comprises three HS6ST isoforms (HS6ST1-3) and seven HS3ST isoforms (HS3ST1, HS3ST2, HS3ST3a/3b, HS3ST4-6)^20^. HS6ST2 ditfers from HS6ST1 and HS6ST3 in its marked preference for HS substrates already N- and 2-O-sulfated^61,62^. HS3ST5, unlike HS3ST2, HS3ST4 and HS3ST6, preferentially modifies HS domains containing pre-existing 6-O-sulfated glucosamine residues^22,63^. The combinatorial specificity conferred by the co-enrichment of HS6ST2 and HS3ST5 at the rat NMJ therefore appears to arise from these sequential substrate preferences, which favor highly sulfated sequences carrying the 2SNS6S3S motifs detected by Hs4c3. In mice, HS3ST3a and HS3ST3b generate products less dependent than HS3ST5 on 6-O-sulfated substrates^20,22^. Rather than relying on several HS3ST3 enzymes, the rat NMJ appears to use HS3ST5 preferentially, allowing one enzyme to generate both gD- and AT-type motifs. This species-specific ditference in HS3ST isoform usage may represent an alternative route to a functionally conserved mode of 3-O-sulfation.

A closer analysis of the structural basis of this potentiating activity showed that it does not depend on the nature of the uronic acids flanking the 3-O-sulfated glucosamine, since both iduronic and glucuronic acids support potentiation. The 2-O-sulfated residues adjacent to the GlcNS6S3S motif are nonetheless required, since their absence enhanced agrin-LRP4 binding yet failed to induce AChR clustering. Enhanced agrin-LRP4 binding is therefore not sutficient on its own to trigger downstream AChR clustering. Specific HS motifs may instead regulate steps downstream of receptor engagement, such as receptor conformational changes, stabilization of signaling complexes, or recruitment of etfectors required for MuSK activation. More broadly, these findings expose the limits of inferring HS function from *in vitro* ligand binding alone. Glycan microarrays and related biochemical approaches have been instrumental in defining protein-binding preferences for sulfated HS sequences^64^, but they do not recapitulate the spatial organization, multivalency, membrane presentation, and proteoglycan context of the cell surface. Sequence requirements for HS biological activity therefore cannot be inferred from binding atfinity and must be assessed in physiologically relevant cellular systems.

### *In vivo HsCst2* and *Hs3st5* co-expression prevents AChR cluster fragmentation in DMD

The cause of AChR cluster fragmentation at the DMD NMJ remains unclear. Some studies describe disrupted NMJ morphology as a direct consequence of muscle fiber degeneration and regeneration^65,66^, while others report fragmentation independent of these processes^10,67^. *HsCst2*/*Hs3st5* overexpression prevents NMJ fragmentation without preventing degeneration and regeneration, so NMJs can be stabilized within an otherwise altered tissue environment. Degeneration and regeneration are therefore not sutficient on their own to drive fragmentation, which must depend on at least one separable mechanism.

AChR clustering at the NMJ is primarily orchestrated by the agrin-LRP4-MuSK signaling pathway¹²˒¹³. Given that gD-type 3S-HS mimics enhance agrin binding to LRP4 and promote agrin-dependent AChR clustering, while having no etfect on AChR clustering on their own *in vitro*, *HsCst2*/*Hs3st5* overexpression may similarly promote AChR aggregation *in vivo* by facilitating agrin and LRP4 recruitment through 3S-HS production. However, we cannot totally rule out the possibility that 3S-HS also modulates other signaling pathways at the NMJ, potentially acting in concert with agrin. HSPGs modulate numerous extracellular signaling systems, including neuregulin/ERBB, Wnt, FGF, and BMP pathways, all of which contribute to NMJ development, maturation, maintenance or regeneration^68–74^.

The DAPC may provide an additional interface for HS-dependent mechanisms, since it links AChR clusters to the basal lamina and the intracellular cytoskeleton and is disrupted in DMD. Through its interaction with utrophin^75^, dystrophin^76^, rapsyn^77,78^, the dystroglycan complex contributes to NMJ organization and stability. Moreover *α*-dystroglycan also binds agrin^79^ and laminins^80^ both involved in AChR clustering^81^. Notably, HS6ST2 co-immunoprecipitated with *β*-dystroglycan from mdx mice overexpressing *Galgt2*^82^, although the significance of this interaction remains unclear. This finding raises the possibility that HS-dependent mechanisms are functionally linked to the DAPC and other components of the NMJ extracellular matrix.

Activation of the agrin-LRP4-MuSK pathway has been proposed as a therapeutic strategy to mitigate DMD^14^. Our results align with that proposal, since co-expression of *HsCst2* and *Hs3st5* preserved NMJ morphology with etfects comparable to, or greater than, those reported after LRP4 overexpression in mdx mice^14^. Rather than targeting the pathway directly, remodeling HS fine structure may otfer an alternative route to stabilizing the postsynaptic apparatus. This is consistent with the role of 3S-HS in NMJ stabilization in *C. elegans*^32^, although identifying the molecules shared between vertebrates and invertebrates will require further work. The etfects of co-expression extended beyond morphological changes to include improved muscle strength, despite the absence of detectable improvements in other established markers of muscle pathology, including centrally nucleated regenerating fibers and fibrosis. Therefore, a parsimonious explanation for our results is that the increase in muscle strength is attributable to NMJs stabilization by 3S-HS rather than other mechanisms. Our results also highlight several avenues for further investigation, which will be addressed in future studies: i) extending treatment duration to maximize the etfects of *HsCst2*/*Hs3st5* co-transduction; ii) directly evaluating the therapeutic potential of 3S-HS mimics *in vivo*; and iii) refining functional readouts to enable a more comprehensive assessment of NMJ integrity.

Finally, 3S-HS may play an increasingly important role in NMJ organization during aging, as their abundance at the NMJ in WT rats increases significantly only at advanced stages of adulthood. This increase may reflect an adaptive response that progressively reinforces NMJ stability in the face of cumulative age-related muscle stress. The physiological relevance of this mechanism in WT animals and its contribution to the aging process remain to be established. Addressing these questions will require further investigation using approaches that specifically inhibit 3S-HS production or function.

### Further implications of 3S-HS engineering for neuromuscular diseases

Beyond DMD, these findings broaden the functional significance of the HS sulfation code and suggest that modulating tissue-specific glycan signatures may otfer a general means of regulating extracellular signaling. NMJ destabilization is a common pathogenic feature of numerous neuromuscular disorders, including spinal muscular atrophy, amyotrophic lateral sclerosis, congenital myasthenic syndromes, and peripheral neuropathies^83–86^. Activation of the agrin-LRP4-MuSK pathway preserves synaptic integrity and delays denervation independently of the underlying genetic defect. Our work establishes the NMJ as a specialized HS microdomain where 3S-HS motifs regulate synaptic stability and provides proof of concept that engineering the HS sulfation code to potentiate agrin-dependent AChR clustering can preserve NMJ function. Because this approach targets the extracellular matrix rather than intracellular pathways, it may apply broadly across genetic disorders atfecting the NMJ.

## Methods

### Animals

The Del 52 rat model was previously generated by generating a 188-bp deletion encompassing exon 52 and introducing a premature stop codon in exon 53 of the dystrophin (*Dmd*) gene^33^. The R-DMDDup 10-17 rat model was established by duplicating exons 10-17 of *Dmd* gene^34^. Both lines were maintained in the Sprague Dawley (SD) rat strain genetic background (Janvier Labs). Wild-type and DMD rats were littermates and were housed in a pathogen-free facility under environmental conditions compliant with European directive 2010/63/EU. Given the X-linked recessive inheritance of DMD, only male rats were included for the experiments. All the procedures including animal handling were validated by the ethics committee of the French ministry (APAFIS#58107-Committee No. 16).

### Tissue clearing and NMJ morphological analysis

Tibialis anterior (TA) muscles from 2- and 7-month-old wild-type, Del 52 and Dup 10-17 SD rats were dissected and immersion-fixed in 4% paraformaldehyde (PFA) in 1× PBS for 3 h at room temperature. Muscles were cut longitudinally into 1-mm-thick slices and post-fixed in 4% PFA. After washing in PBS, slices were stored at 4°C in PBS containing 0.2% sodium azide. For immunolabeling, slices were permeabilized in PBST containing 0.2% Triton X-100 for 48 h at room temperature and then incubated for 6 days at 37 °C in dilution butfer (1% normal goat serum, 0.2% Triton X-100 and 0.2% sodium azide in PBS, 0.1 M) supplemented with anti-SV2 antibody (DSHB, AB_2315387; 1:100) and anti-neurofilament antibody (DSHB, AB_531793; 1:100). After washing in washing butfer (PBST supplemented with 3% NaCl) slices were incubated for 4 days at 37°C in the dark with agitation with Alexa Fluor 488 goat anti-mouse IgG1 (Thermo Fisher, A-21121; 1:500) and Alexa Fluor 647-conjugated *α*-bungarotoxin (Thermo Fisher, B35450; 1:500) and nuclei counterstained with DAPI. For clearing, slices were incubated in pre-warmed RapidClear 1.52 (Sunjin Lab, RC152001) for 24 h at 25°C in the dark. Samples were mounted in RapidClear medium using a 1-mm spacer to maintain sample thickness and prevent bubble formation. Images were acquired using a Zeiss spinning-disk confocal microscope equipped with a Yokogawa CSU-X1 scanning unit and a 25× water-immersion objective (numerical aperture 1.51). Image stacks were acquired at a voxel size of 0.26 × 0.26 × 1.0 µm using identical acquisition settings for all experimental groups. The camera, excitation and emission settings, laser power, gain, stack depth, and software version were identical for all experimental groups. Image stacks were processed in Fiji (ImageJ; version 2.14.01.54f) by merging channels and defining a standardized three-dimensional region of interest encompassing the complete NMJ. Thirty NMJs from three animals per condition and fifteen NMJs from five animals per condition were analyzed for NMJ morphological characterization in DMD and for the *in vivo* transduction experiments, respectively. Processed image stacks were imported into Imaris (version 10.2.0; Oxford Instruments) for three-dimensional reconstruction. Axo-presynaptic structures, defined by neurofilament and SV2 immunoreactivity, and postsynaptic structures, defined by *α*-bungarotoxin labeling, were segmented separately. Innervation was scored manually according to morphological continuity between presynaptic and postsynaptic channels as follows: 0, non-innervated; 1, partially innervated; and 2, fully innervated. Colocalization was quantified using the Imaris Overlap Volume Ratio statistic. Volume, surface area, and contact parameters were extracted using the Imaris Statistics module and exported for statistical analysis. Thresholding and segmentation parameters were defined separately according to prespecified criteria.

### Tissue preparation for immunofluorescence

Tibialis anterior, diaphragm and soleus muscles were dissected after animal terminal euthanasia. The fascia was removed, and tissues were oriented transversely, for cross-section analysis, or longitudinally, for NMJ analysis, on cork supports using tragacanth gum, snap-frozen in isopentane rapidly cooled by liquid nitrogen and stored at -80°C. Ten-micrometer cryosections were prepared, air-dried and fixed in methanol at -20°C. Sections were then washed in PBS, permeabilized in 0.1% Triton X-100 in PBS for 25 min at room temperature, washed and blocked in PBS containing 10% BSA for 30 min. Sections were incubated overnight at 4°C in the dark with the following primary antibodies. For NMJ analysis, Hs4c3^87^ (1:50), mouse Lrp4 (NeuroMab, clone N207/27; 1:500), rabbit Musk (Merck, ABS549; 1:500), mouse MF20 (DSHB AB_2147781; 1:300). For gain-of-function analysis, chicken GFP (Abcam, EPR140104; 1:1000), rabbit RFP (Tebubio, 039600-401-379; 1:500) rabbit laminin (Sigma, L9393; 1:500), mouse collagen VI (Merck, MAB1944; 1:1000). Sections were washed in PBS and incubated with the corresponding Thermo Fisher secondary antibodies, Alexa Fluor 488 goat anti-mouse (Thermo Fisher, A-1100; 1:500), Alexa Fluor 555 goat anti-rabbit (Thermo Fisher, A-21429; 1:500) or Alexa Fluor 488 goat anti-chicken (Thermo Fisher, A-11039; 1:500) for 60 min at room temperature. Rabbit anti-VSV (Thermo Fisher, PA1-30138; 1:100) followed by goat anti-rabbit IgG A555 was used for Hs4c3 Ab recognition. Sections were counterstained with Alexa Fluor 647-conjugated *α*-bungarotoxin (Thermo Fisher, B35450; 1:500) and DAPI (1:500). Fluorescence intensity at NMJ was quantified using a mask generated from *α*-bungarotoxin staining to delineate NMJ surfaces. Mean fluorescence intensity was measured for 70 NMJs. Background was measured in tissue immediately adjacent to NMJ and subtracted from the NMJ intensity value before quantification. Acquisition settings were kept constant across experimental groups. Signal intensity was normalized to mean *α*-bungarotoxin intensity to estimate NMJ labeling relative to cholinergic-receptor signal. For statistical analysis, NMJ measurements were averaged per animal.

### Plasmid construction

Expression constructs were generated using a modified pCIG backbone containing a CMV enhancer upstream of a CAAG promoter, an internal ribosome entry site (IRES) followed by a nuclear-localized GFP reporter and bacterial selection elements (illustrated in Fig. S6). A multiple-cloning site containing EcoRV and XmaI sites was used for insertion of the target coding sequences. Coding sequences for Hs6st2 and Hs3st5 were cloned to generate pCMV/CAAG-Hs6st2 and pCMV/CAAG-Hs3st5, respectively. Constructs were verified by restriction digestion and Sanger sequencing and were prepared using an endotoxin-free plasmid purification method. Plasmid maps were generated in ApE v3.1.6. The bacterial strain used for plasmid amplification was competent DH10B E. coli (Thermo Fisher, EC0113), and purification was performed using NucleoSpin Plasmid EasyPure and NucleoBond Xtra Midi kits (Macherey-Nagel, 740727.50 and 740410.50).

### NeoPG mimics synthesis

Procedures for neo-proteoglycan synthesis have been reported previously^44,45,88^. 7-mer peptide backbones were synthesized through manual solid-phase peptide synthesis on Rink amide resin. The peptide backbone was tagged with Cy5 dye at the N-terminal for real-time imaging and conjugated to Cholesterol-DBCO at the C-terminal to immobilize on the cell surface. The obtained 7-mer peptides were further conjugated to respective carbohydrate moieties employing SPAAC reaction. Sugars M14 to M18 were conjugated to 7-mer peptides to obtain the corresponding proteoglycans. The synthesis and characterization of the neoPGs M14, M15, M16, M17 and M18 were previously reported. Briefly, the sugar derivatives were dissolved in anhydrous DMF and stirred for 5 min in the presence of Et3N, followed by dropwise addition of DBCO-NHS solution in anhydrous DMF. The reaction mixture was stirred for 12 h at room temperature. Upon purification, the obtained product (1.2 equiv. per azide on the peptide) was dissolved in H2O and added to a solution of peptide with azide linker (1 equiv.) dissolved in CHCl_3_: MeOH (1:1). The reaction mixture was stirred for 7-8 days, evaporated and filtered through a 10-kDa cutotf filter to obtain the final neoPGs.

### C2C12 cell culture and AChR clustering assay

C2C12 myoblasts were grown on a collagen I coating prepared at a final concentration of 50 µg/ml by diluting 2.5 ml of collagen I stock solution (3 mg/ml) (Gibco, A10483-01) in 147.5 ml of 20 mM acetic acid, yielding a final volume of 150 ml. C2C12 were seeded on collagen I-coated chamber slides at 15,000 cells/cm² in proliferation medium consisting of DMEM (Gibco, 41966-029) supplemented with 15% fetal bovine serum (Eurobio, CVFSVF00), 1% penicillin-streptomycin (Gibco, 15140-122) and 1% L-glutamine (200 mM 100X stock; Gibco, 25030-024). Cells were maintained for 48 h, with a medium change after 24 h. At confluency, cells were washed with PBS, and the proliferation medium was replaced with ditferentiation medium (changed twice at 48-hour intervals) consisting of DMEM supplemented with 2% horse serum (Gibco, 16050-122). On the fifth day of ditferentiation, recombinant rat agrin (R&D Systems, 550-AG/CF) was added to the cell cultures at a final concentration of 3 ng/ml (approximately 30 pM) for 16 h; the cells were then washed with PBS and fixed for analysis. C2C12 myotubes were routinely used for agrin-induced AChR clustering assays, involving AChR labeling with *α*-bungarotoxin, and were also subjected to immunostaining using the antibodies employed for tissue sections, as previously described. Spontaneous and agrin-induced AChR clusters were identified in a size range from 1 µm to 100 µm and quantified per well. The number of AChR clusters was normalized to the number of fused nuclei, defined as nuclei associated with MYH-positive myotubes. Images were acquired from randomly selected fields by an investigator blinded to experimental conditions. Each condition was analyzed in either 3 or 6 independent experiments, depending on the experiments, with two replicate wells per experiment, each totaling 300 to 600 fused nuclei analyzed.

### C2C12 cell specific treatments

For transfection experiments involving plasmids encoding the sulfotransferases Hs6st2 and Hs3st5, C2C12 cells were seeded and cultured for 24 h prior to transfection. Endotoxin-free plasmid DNA was diluted to the required concentration. For single transfections, 750 ng of DNA was used per well. For co-transfections, 375 ng of each plasmid was used, resulting in a total of 750 ng of DNA per well. Transfection complexes were prepared using Lipofectamine 3000 (Invitrogen L3000001) according to the manufacturer’s instructions. Lipid and DNA mixtures were prepared separately, combined, and incubated for 15 min at room temperature before being added to the cells. Finally, the cells were treated with rat agrin recombinant protein (R&D Systems) 6 days post-transfection for the agrin-induced AChR clustering assay. For competition experiments, HS-specific 12-unit oligosaccharides (12-mers) were added simultaneously with neuronal agrin to the transfected or non-transfected cell cultures. Three 12-unit oligosaccharides (Glycan Therapeutics) with specific sulfation profiles were used (schematized in Fig. 5a)^23^. The oligosaccharides (stored at 100 µg/ml) were diluted to 250 ng/ml immediately before use. Control conditions included a non-3-O-sulfated control oligosaccharide (Oligo CTRL) and a cellular control without Hs6st2/Hs3st5 co-transfection. After 16 h of incubation, the cells were fixed, labeled for AChRs, and immunolabelled for MF20 or GFP. The number and length of AChR clusters were quantified as previously described. Data represent n = 3 to 6 independent experiments per condition. For some experiments, neoPG mimics diluted in PBS were co-administered with agrin at a neoPG concentration of 2 µM.

### ELISA-like assay

Recombinant His-tagged agrin (R&D Systems) was immobilized overnight at 4 °C on anti-His antibody-pre-coated wells of a 96-well microtiter plate (His Tag Antibody Plate, GenScript) by adding 400 ng/well of agrin in 150 µl TBS-0.05% Tween. After washing, wells were incubated overnight at 4 °C with the same amounts of mouse ectoLRP4-AP (as determined by AP activity) present in conditioned medium^46^, in the presence or absence of 2 µM O-sulfated HS mimics, in TBS-0.05% Tween, 1 mM MgCl2. After four washes with TBS-0.05% Tween, 1 mM MgCl2, the amount of AP bound to the wells was measured using p-nitrophenyl phosphate as a substrate (Sigma-Aldrich). The yellow reaction product was read at 405 nm in an Enspire (PerkinElmer) microtiter plate spectrophotometer.

### Transcriptomic analysis

For snRNAseq, diaphragm muscle samples were collected from two 11-month-old wild-type and two Del 52 rats. After removal of the fascia, 50 mg of tissue per animal was dissected from the central band of the muscle, a region enriched in NMJs as described^35^. After pooling the two samples of each condition together, samples were processed to isolate nuclei according to previous protocol^89^. Approximately 100,000 nuclei were collected by fluorescence-activated cell sorting (FACS), and approximately 25,000 nuclei were loaded into one channel of a Chromium X instrument using the Chromium Single Cell 3′ Gene Expression v4 reagent kit (10x Genomics). The number of recovered nuclei was 6,771 for the WT condition and 4,374 for the DMD condition. Following capture and lysis, cDNA was synthesized and amplified by PCR for 11 cycles according to the manufacturer’s protocol. Amplified cDNA was used to generate Illumina sequencing libraries. Two samples were sequenced on a single NextSeq500 Illumina flow cell in paired-end mode (Read 1: 28 cycles; Read 2: 54 cycles; Index 1: 10 cycles). Raw sequencing data were processed using Cell Ranger v8.0.0 (10x Genomics), which converted base call files into FASTǪ format, aligned reads to a custom mRatBN7.2 transcriptome using the STAR aligner^90^, and quantified transcript expression per cell using Chromium barcodes. Sequencing data were demultiplexed into FASTǪ files using the mkfastq command in Cell Ranger v8.0.0 (10x Genomics). A gene-by-nucleus count matrix was generated for each sample. Low-quality nuclei were defined as nuclei with fewer than 200 detected features and mitochondrial transcript content greater than 5%. Features detected in fewer than three nuclei were removed. Nuclei and features were filtered independently for each sample using Seurat v5.4.0. Samples were merged into a single Seurat object and normalized using the LogNormalize method with a scale factor of 10,000. The 2,000 most highly variable features were identified using the variance-stabilizing transformation method. Data were scaled and centered, followed by principal-component analysis. Batch etfect correction was performed by integrating the datasets through canonical correlation analysis. UMAP dimensionality reduction and construction of a shared-nearest-neighbor graph were performed using R v4.3.3. Clustering was performed using the Louvain algorithm at a resolution of 0.5, yielding 15 clusters. Clusters were manually annotated using marker genes. Ditferential-expression analysis was performed using the FindMarkers function in Seurat v5.4.0 with the Wilcoxon rank-sum test. Multiple-testing correction was performed using the Benjamini-Hochberg method. Genes with an adjusted p-value < 0.05 were considered ditferentially expressed. Over-representation analysis (ORA) was performed to identify enriched Gene Ontology (GO) terms using the clusterProfiler R package. The analysis was conducted using gene symbols as identifiers against the rat genome annotation database. Functional enrichment was assessed across the Biological Process, Molecular Function and Cellular component categories. Statistical significance was determined using the hypergeometric test, with p-values adjusted for multiple testing via the Benjamini-Hochberg method. Terms with an adjusted p-value < 0.05 were considered significantly enriched

For RT-qPCR, RNA was extracted from C2C12 cells collected at confluence and 5 days after induction of ditferentiation. RNA was extracted using NucleoSpin RNA XS (Macherey-Nagel, 740902.05), and RNA quantity was assessed by spectrophotometry. Reverse transcription was conducted using a Biometra TOne thermocycler and the reagents listed in Appendix 1. The resulting cDNA was used for quantitative real-time PCR (RT-qPCR) analysis. For each sample, cDNA corresponding to 25 ng of input RNA was used for quantitative real-time PCR. The primer sequences of the mouse heparan sulfate sulfotransferases have been previously published^91^. *Gapdh* (Primer Forward: GACGGCCGCATCTTCTTGT, Reverse: CACACCGACCTTCACCATTTT) was used as the reference gene after confirming its stability across experimental groups. No-template controls were prepared by replacing cDNA with nuclease-free water. Each sample was measured in three technical replicates. Relative expression was calculated using the ΔCT method.

### AAV-mediated *HsCst2* and *Hs3st5* transduction

Recombinant muscle-targeted MyoAAV2a vectors were used to drive transgene expression under the control of the muscle-specific MHCK7 promoter. Two vectors were generated: MHCK7-*HsCst2*-T2A-nls-*mCherry* and MHCK7-*Hs3st5*-T2A-nls-*GFP*. A Kozak consensus sequence was placed upstream of each transgene start codon. Downstream of the *HsCst2* and *Hs3st5* coding sequences, the *Thosea Asigna* virus 2A-like (T2A) self-cleaving sequence linked the transgenes to nuclear-localized mCherry and GFP reporters, respectively, enabling coordinated expression from a single transcript^92^. Both vectors also contained a shortened woodchuck hepatitis virus post-transcriptional regulatory element (WPRE3) and a Simian virus 40 (SV40) late polyadenylation signal^93^. Coding sequences were codon-optimized using the CUSTOM server (CRG) as described^94^. Designed sequences and MyoAAV2a vectors were obtained from PackGene Biotech (China), via transient transfection of HEK293 cells, purified by atfinity chromatography, and released after quality control comprising SYBR Green qPCR genome titration, reported as 4 × 10¹² GC/ml; SDS-PAGE purity assessment, which showed no additional bands beyond the expected pattern; and endotoxin testing by Limulus amebocyte lysate (LAL) assay at <10 EU/ml. Aliquots were stored at -80°C and thawed on ice immediately before use. MyoAAV vectors were injected into the tibialis anterior of two-month-old male Del 52 and wild-type rats on an SD background. On the day of injection, buprenorphine (30 µg/kg, subcutaneous) was administered 20 min before anesthesia. Animals were anesthetized with isoflurane (4% for induction and 2% for maintenance) and placed on a heating pad. Ocular lubricant (Viskyal) was applied to prevent corneal drying. Intramuscular injections were performed using a 30-gauge needle and Hamilton syringe. Each muscle received 100 µl of viral suspension containing 3 × 10^11^ vg. The experimental group received both vectors, with the intended dose divided equally between vectors (1.5 × 10^11^ vg of each vector). Control animals received the control MyoAAV construct MHCK7-nls-GFP at 3 × 10^11^ vg. Animals recovered in a warmed cage at 28 °C with supplemental oxygen and infrared heating for at least 30 min, until spontaneous locomotion resumed, then rats were returned to their home cages and housed under standard conditions. Animals were inspected for general health and injection-related adverse etfects 6 h after injection and daily thereafter, in accordance with institutional animal-care guidelines.

### Muscle force measurement and NMJ morphological analysis

Three months after injection, muscle force was assessed before sacrifice using an isometric force transducer coupled to a stimulator for sciatic-nerve activation. Because this procedure required deeper anesthesia, buprenorphine (50 µg/kg, subcutaneous) was administered 20 min before ketamine (50 mg/kg, subcutaneous) and medetomidine (0.4 mg/kg, subcutaneous). Animals were continuously monitored for body temperature, respiratory rate and electrocardiogram when required. Oxygen-enriched air was provided throughout the procedure to reduce the risk of hypoxemia and respiratory arrest. Following induction of deep anesthesia, the skin over one hindlimb was incised longitudinally in the craniocaudal direction to expose the underlying musculature. The distal tendon of the tibialis anterior was transected and secured with silk suture to the lever arm of a dual-mode force transducer (Aurora Scientific, model 305B). The foot was blocked, and the knee was firmly immobilized using a bone fixation device. The sciatic nerve was exposed through a lateral incision in the thigh and stimulated with a bipolar hook electrode using supramaximal square-wave pulses (0.1 ms) to elicit muscle contractions with a train duration of 500 ms. Supramaximal current intensity was first established in constant-current mode using 100-Hz tetanic trains (500 ms duration). Optimal muscle length (L0) was determined by adjusting muscle length until maximal tetanic force was achieved during stimulation at 100 Hz. After a 1-min rest period, peak absolute force was measured during two tetanic stimulation protocols: indirect stimulation of the sciatic nerve in constant-current mode followed by direct stimulation of the muscle of interest in constant-voltage mode. A 1-min rest period was applied between the two stimulations to prevent any neuromuscular fatigue. In both conditions, the stimulation intensity was set at a supramaximal level (corresponding to 120% of the minimum intensity required to reach the force plateau of a fused tetanus). Force signals were acquired using Dynamic Muscle Control software and analyzed with Dynamic Muscle Analysis (Aurora Scientific). Specific maximal force was calculated by dividing peak absolute force (N) by muscle mass (g) and is reported as N/g. At the end of the procedure, rats were euthanized by exposure to carbon dioxide (CO2) in accordance with institutional and regulatory guidelines.

### Statistical analysis

Statistical analyses were performed using Prism 9.5.1 (GraphPad). Unless otherwise specified in the corresponding figure legend, all group comparisons were performed with the non-parametric Kruskal-Wallis test followed by Dunn’s test for multiple comparisons. The definition of the biological replicate and the number of independent experiments are reported in each figure legend, as is any test that departs from this default. Data are presented as mean ± s.e.m. All tests were two-sided, and p < 0.05 was considered statistically significant. For measurements nested within animals, animal identity was treated as the experimental unit.

## Data, code, and materials availability

Raw and processed snRNA-seq data are accessible in [repository] under accession number [accession]. Analysis code will be made available at [repository or URL]. Plasmid sequences, vector maps, and other materials required to reproduce the experiments will be available from the corresponding author upon reasonable request.

## Acknowledgments

We thank Laurent Guillaud for genotyping of animals, Kaouthar Kefi, Juliette Berthier and Baptiste Periou for technical advice, Johanna Abrigo-Leon for initially helping in intramuscular AAV injection and in the process of obtaining authorization for the project from the French Ministry of Higher Education, Research and Space, and Alina Blédéa for daily zootechnical monitoring; we thank Sassi De Laat for IT analysis consulting; F.D. particularly thanks Alwyn Dady & Isabella Scionti for scientific discussion during the thesis process. We thank the “Institut Clinique de la Souris” -PHENOMIN (Illkirch, France) for the establishment of the Del 52 and Dup 10-17 rat mutant lines; we acknowledge the ImagoSeine core facility of the Institut Jacques Monod, member of the France BioImaging infrastructure (https://ror.org/01y7vt929) supported by the French National Research Agency (ANR-24-INBS-0005 FBI BIOGEN) and GIS-IBiSA. This work was supported by funding to F.R. from the “Association Française contre les Myopathies” (AFM) via TRANSLAMUSCLE I, II and III programs (projects 19507, 22946 and 30138), the Astellas Gene Therapies grant AT753-201, and the “Fondation pour la recherche médicale” (FRM, grants EǪU20200301021 and SPF20170938733), the Labex REVIVE (1RGE12111PESPESA01) and European MAGIC Project consortium grant101080690. F.D. was awarded a PhD scholarship from the UPEC Doctoral School of Life and Health Sciences. A.C. was awarded a Raman-Charpak postdoctoral fellowship to work in O.S.’s group. O.S. extends warm thanks to Aymeline Vandestienne and Sylvie Manin for their ongoing assistance in organizing the team’s work.

## Authors contributions

Conceptualization: O.S. conceived, developed, and led the project; Methodology: O.S., F.D. Investigations: F.D. performed Cell Culture, Histology, Imaging, most in vitro/in vivo experiments and analysis; F.D. and F.C. performed snRNAseq; S.B.R. and O.S. performed transcriptomic analysis; F.A. and F.D. performed molecular biology; R.K., A.C. and S.A. performed HS mimic development and synthesis; A.D. developed and performed LRP4 ELISA assay; F.R. and L.T. developed and provided DMD models; L.T. established all anesthesia, analgesia, and clinical monitoring protocols for the DMD rats in experimental procedures. A.R. contributed to preliminary discussions on the control of muscle cells by the nervous system. M.G.T. developed the original protocol of tissue transparency. M.L. and F.D. performed muscle stimulations and recording. Writing: O.S. and F.D. wrote the original draft; all authors edited the original version of the manuscript. Funding acquisition: F.R. and contributors (AFM); O.S. and R.K. (A.C. Fellowship).

---

**Figure S1.**
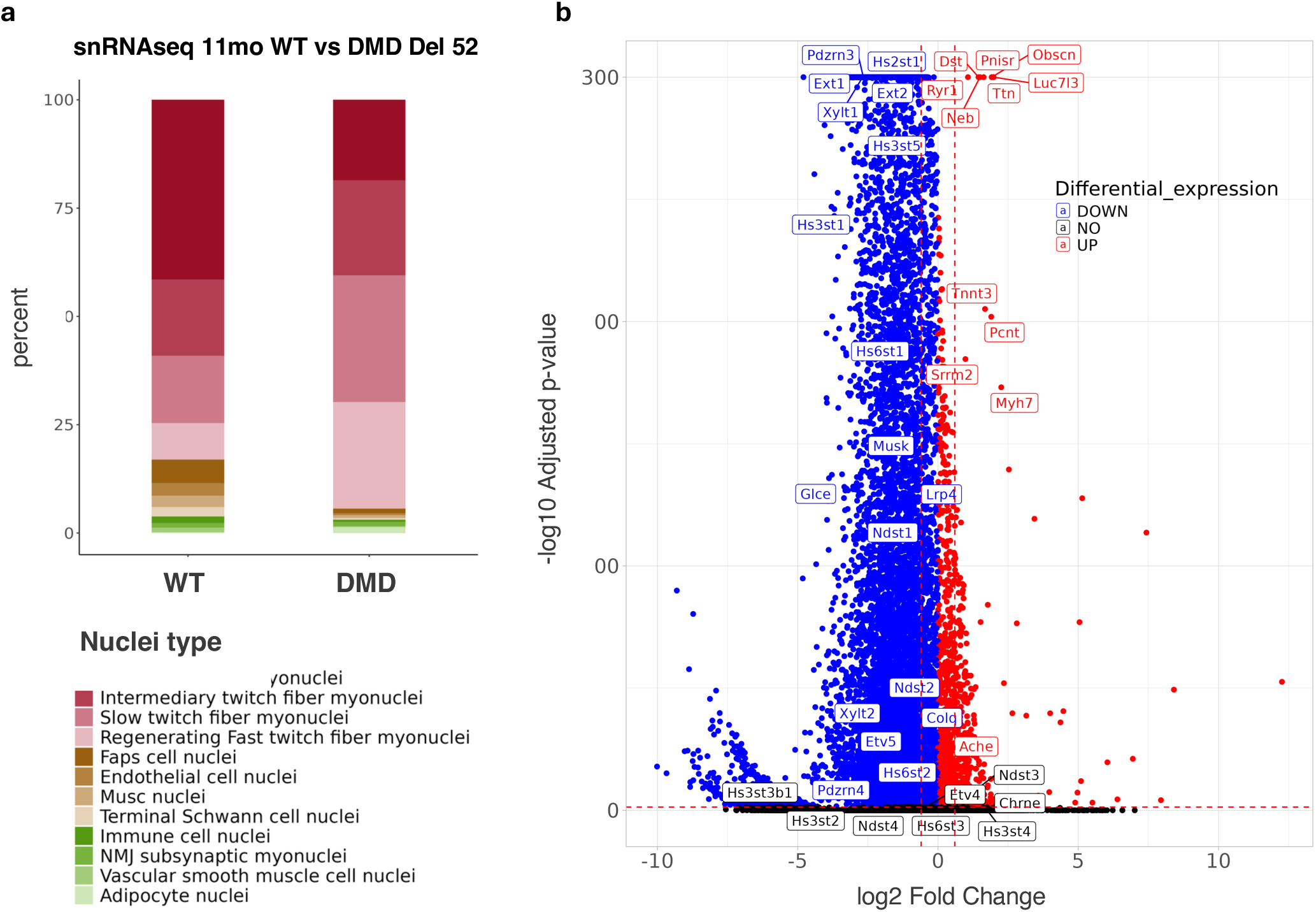
Single-nucleus transcriptomic changes regarding nuclear composition and gene expression in the diaphragm of 11-month-old WT and DMD Del 52 rats. **a**, Stacked bar plot displaying the relative proportion of nuclear populations identified by snRNAseq in WT and DMD diaphragm. **b**, Volcano plot of differentially expressed genes (DEG) in DMD versus WT diaphragm (11,930 DEG over 17,654 genes). Labels highlight heparan sulfate network genes together with top 20 DEG. Coloration is based on differential expressions from the Wilcoxon statistical test.

**Figure S2.**
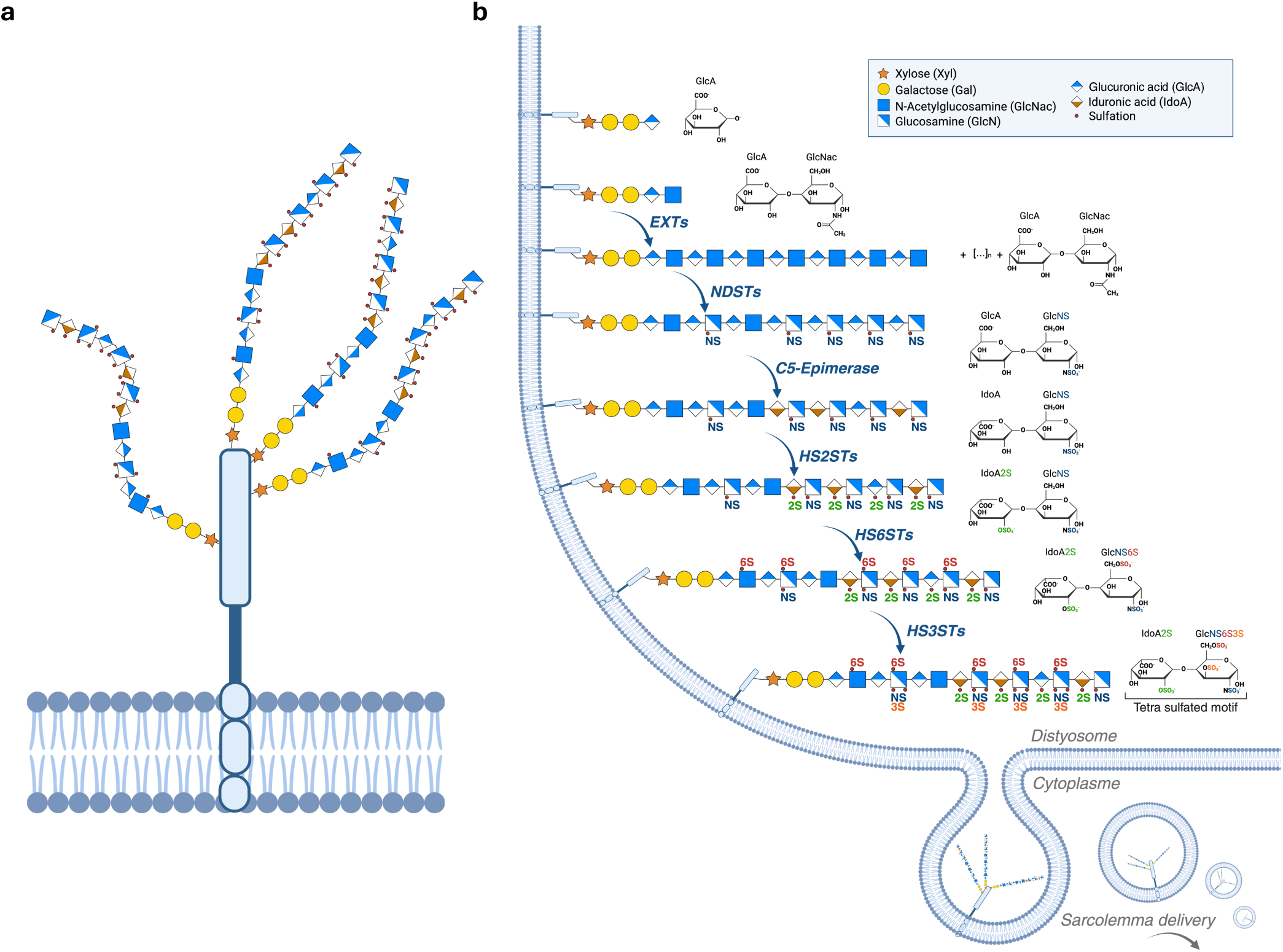
Overview of stepwise synthesis and modifications of heparan glycosaminoglycan chains. **a**, Schematic representation of a membrane-anchored HSPG, showing the core protein associated with heparan sulfate glycosaminoglycan chains extending from the cell surface. b, Stepwise enzymatic biosynthesis and modifications of heparan sulfate chains in the Golgi compartment, beginning with xylose (Xyl), galactose (Gal), and glucuronic acid (GlcA) linkage to the core protein, followed by chain polymerization by EXT enzymes (alternating GlcA and N-acetylglucosamine, GlcNAc), N-deacetylation/N-sulfation by NDSTs (GlcNS), C5-epimerization of GlcA to iduronic acid (IdoA), 2-O-sulfation by HS2STs (2S), 6-O-sulfation by HS6STs (6S), and 3-O-sulfation by HS3STs (3S), generating rare tetrasulfated 3S-HS motif. Mature HSPGs are subsequently trafficked through the Golgi/trans-Golgi network and delivered to the sarcolemma. *Created in BioRender. Relaix, F. (2026)* https://BioRender.com/l0bj1yp.

**Figure S3.**
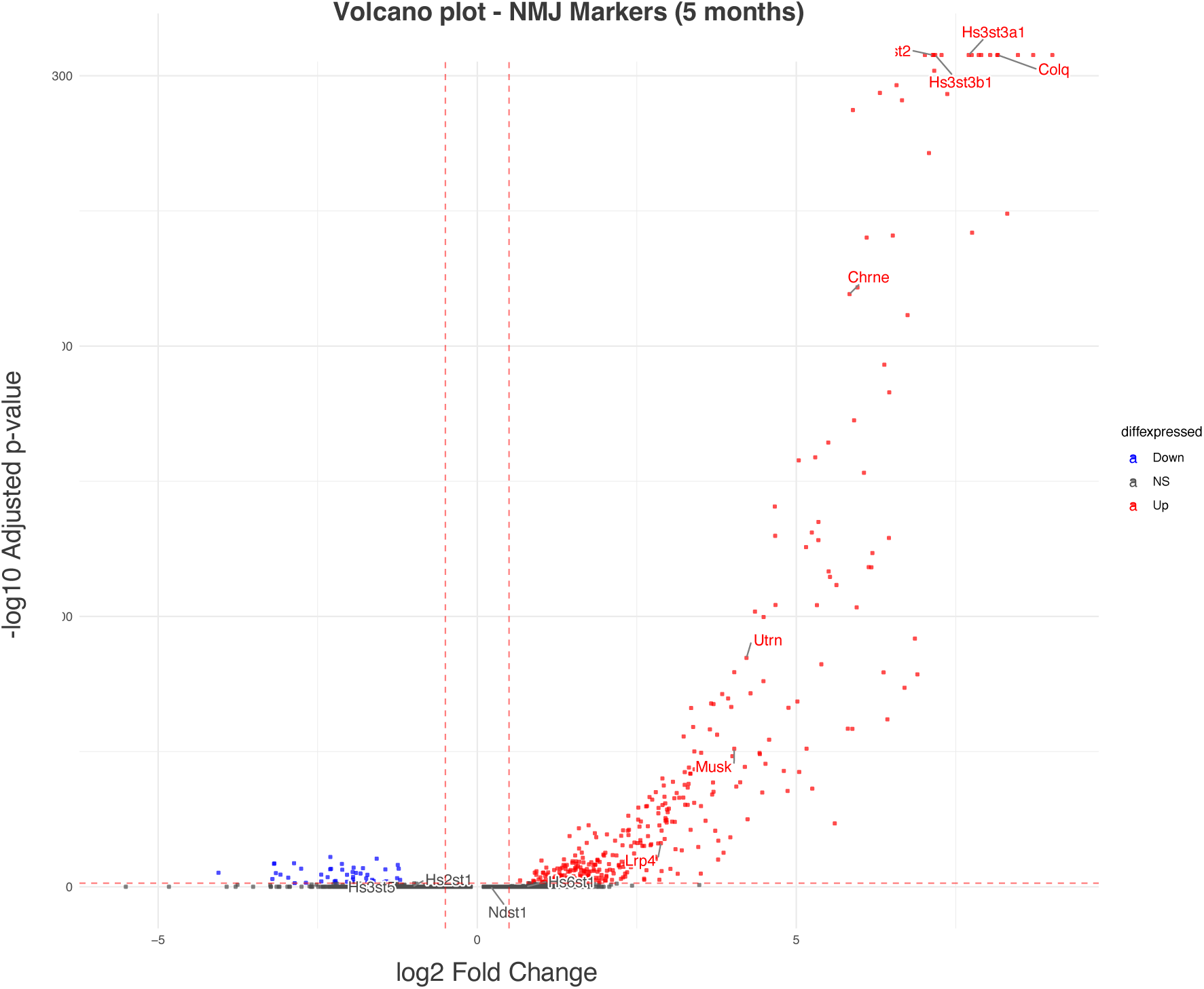
*Hs6st2* and *Hs3st* family members are enriched at the mouse NMJ. Volcano plot of differentially expressed NMJ marker genes in mouse muscle (5 months; MyoAtlas dataset37), comparing NMJ to non-NMJ myonuclei. Established NMJ genes (Colq, Chrne, Musk, Lrp4, Utrn) and heparan sulfate sulfotransferases (Hs6st2, Hs3st3a1, Hs3st3b1, Hs3st5, Hs2st1, Hs6st1, Ndst1) are highlighted. Genes are colored by differential expression status (in blue, downregulated at the NMJ; in red, upregulated at the NMJ; in grey, not significant).

**Figure S4.**
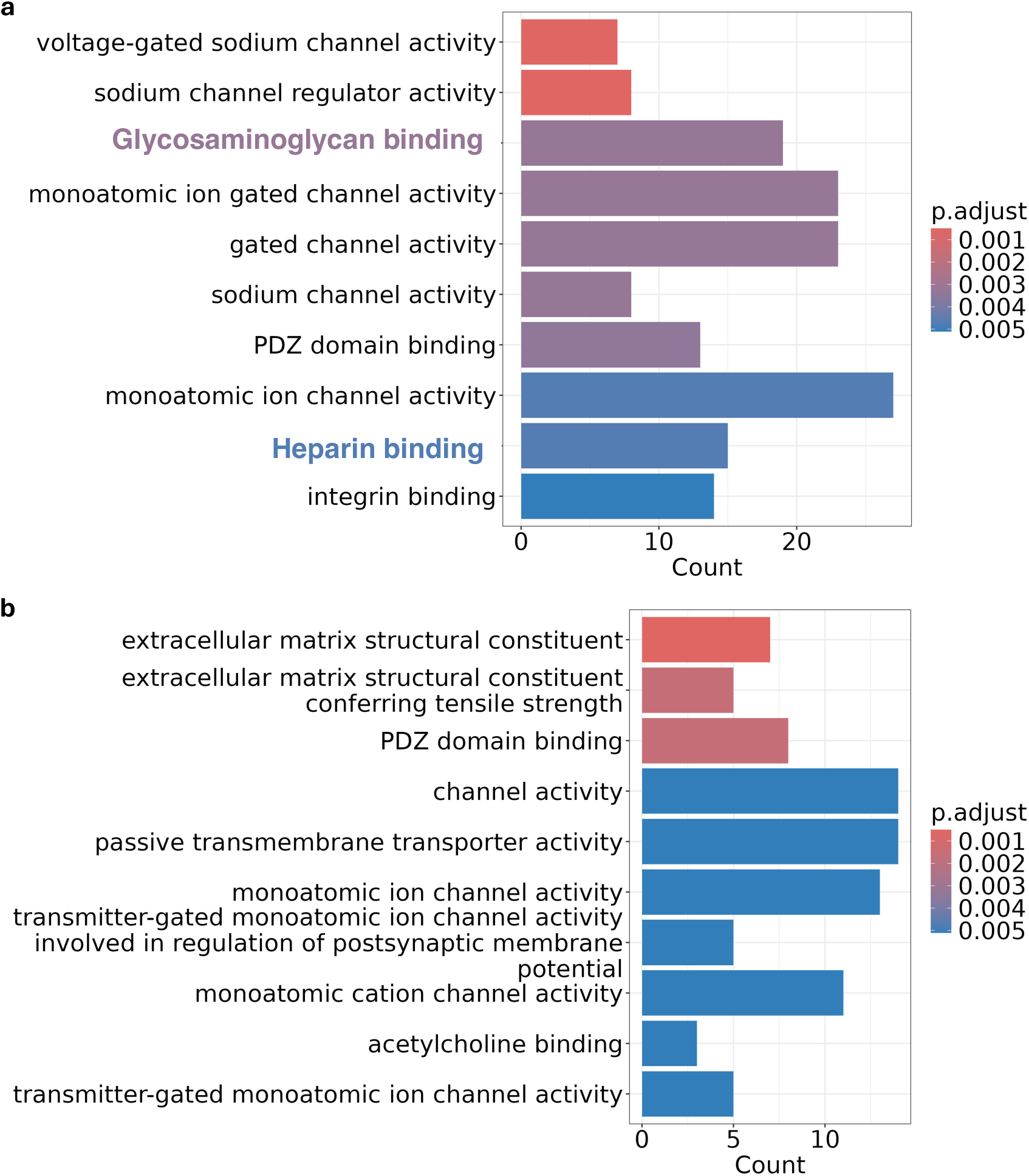
Glycosaminoglycan and heparin binding are enriched molecular functions at the WT but not DMD NMJ. **a**, The top 10 molecular pathways involving genes enriched at the NMJ compared to all clusters in WT (p < 0.05). “Glycosaminoglycan binding” and “Heparin binding” (highlighted) are among the top enriched terms. **b**, Gene Ontology (GO) terms “Glycosaminoglycan binding” and “Heparin binding” are absent from the top 10 molecular pathways involving genes enriched at the NMJ in DMD. ORA (Over-representation analysis) using Gene Ontology database. Colors indicate adjusted p-value (p. adjust).

**Figure S5.**
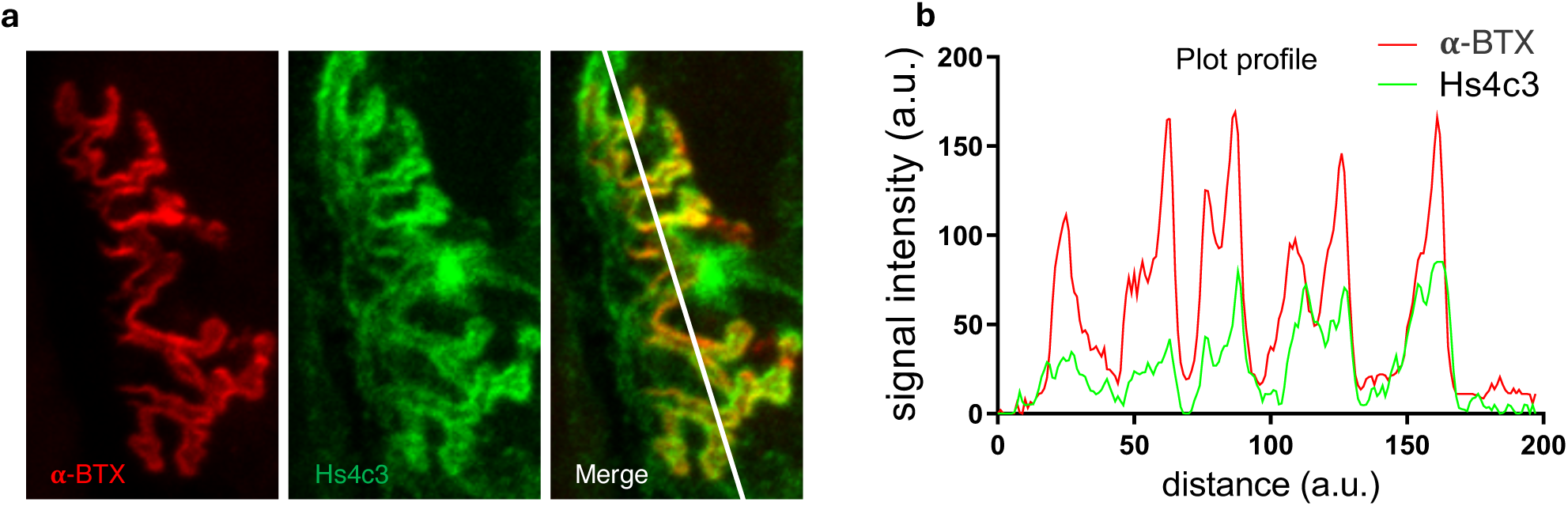
Hs4c3 antibody labeling co-localizes with AChR at the NMJ. a, Representative 5-months-old mouse NMJ labeled for AChR (α-bungarotoxin, in red) and 3S-HS (Hs4c3, in green), with merged image showing the line used for the intensity profile in (b). b, Signal intensity profile of α-bungarotoxin (α-BTX, in red) and Hs4c3 (in green) along the line shown in (a), illustrating colocalization of the two signals at the NMJ.

**Figure S6.**
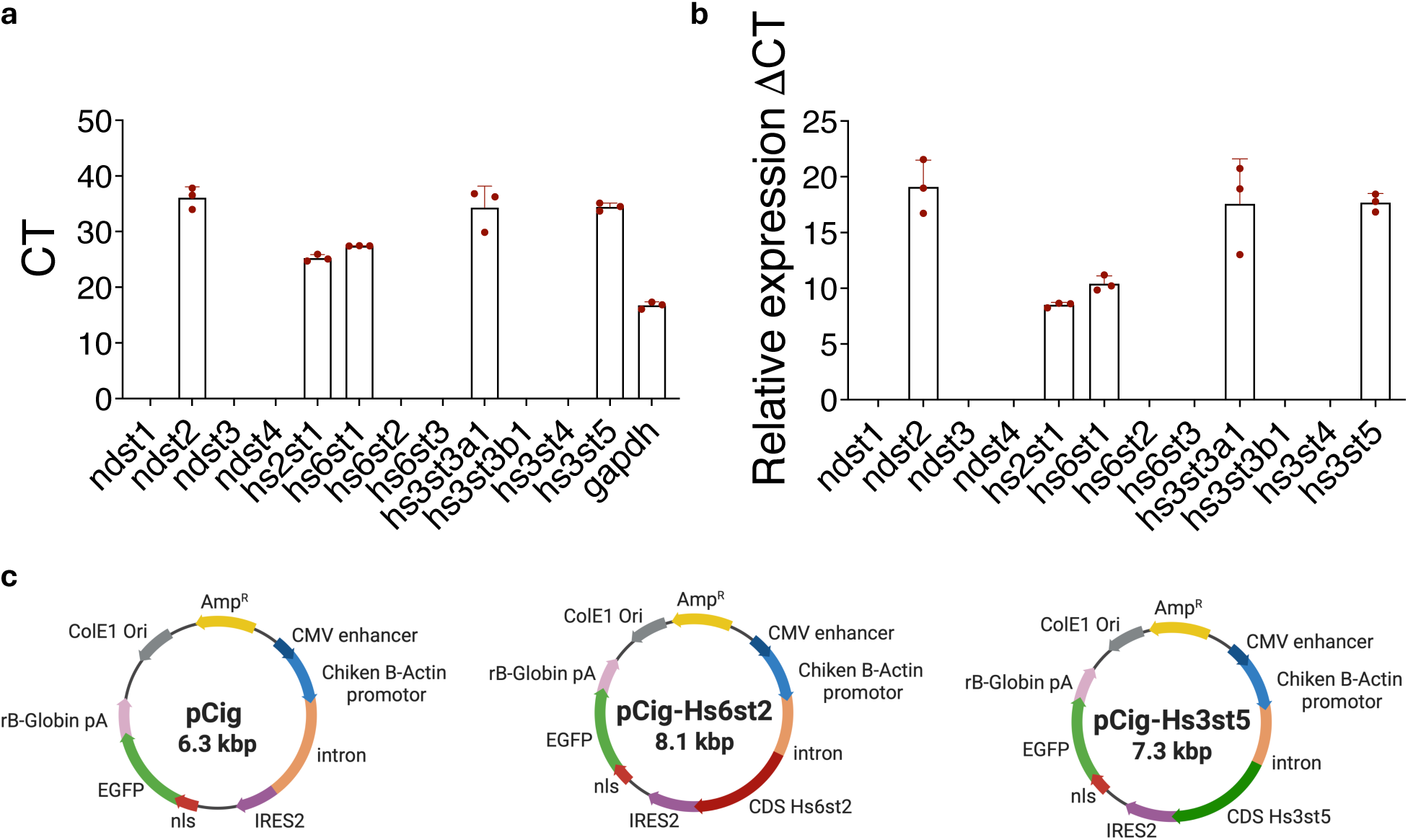
Heparan sulfate sulfotransferase expression profile in C2C12 cells and expression vectors. **a**, Raw expression levels (Ct) of heparan sulfate sulfotransferases in C2C12 myotubes 5 days after differentiation. **b**, Relative expression (ΔCT) of the same transcripts, normalized to glyceraldehyde 3-phosphate dehydrogenase (*Gapdh*). Data are shown as mean ± s.e.m.; each dot represents an independent experiment. **c**, Maps of expression vectors used for gain-offunction experiments: pCIG (control, nlsGFP), pCIG-*Hs6st2* (*Hs6st2*-IRES2-nlsGFP), and pCig-*Hs3st5* (*Hs3st5*-IRES2-nlsGFP).

**Figure S7.**
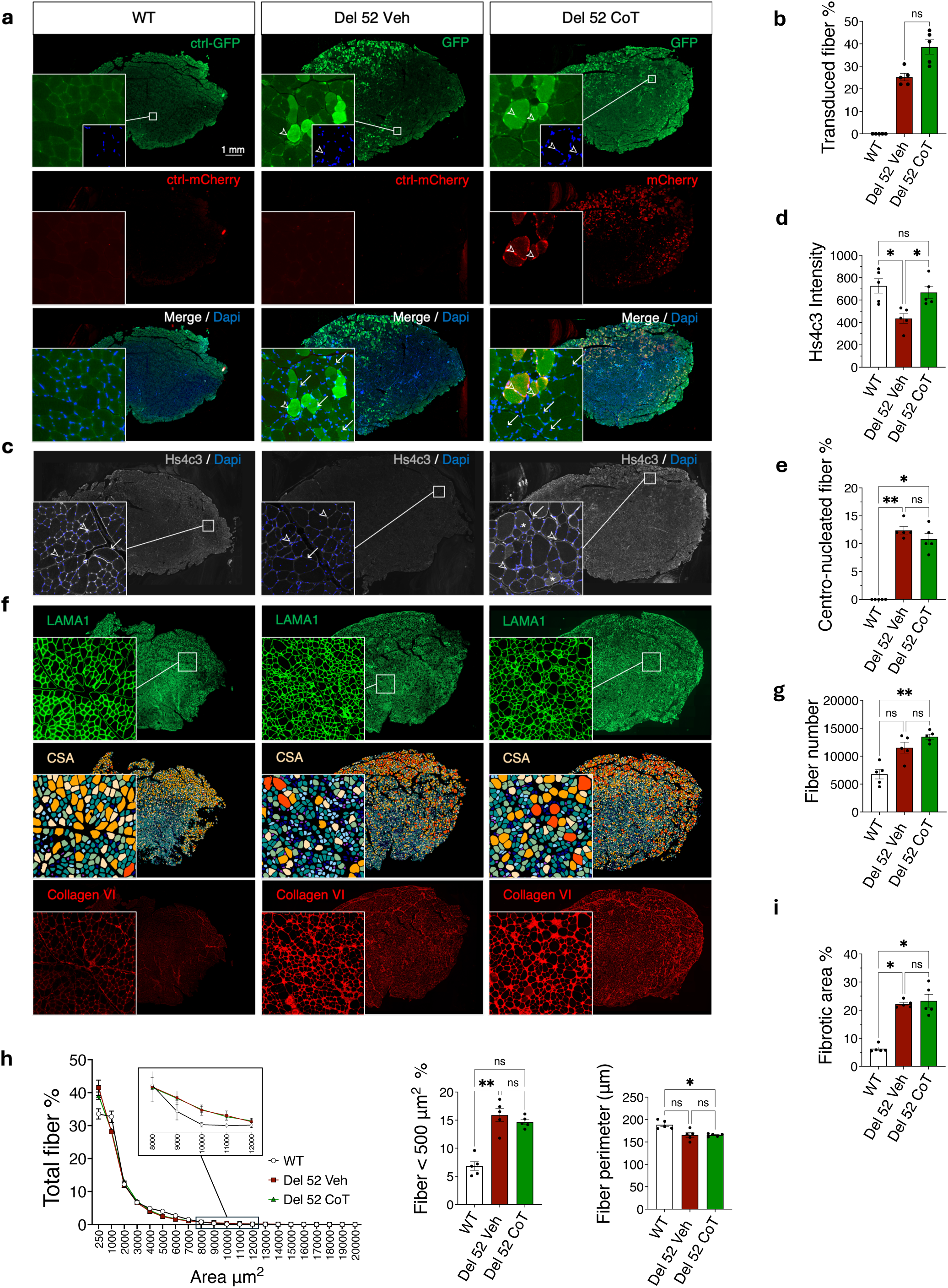
Intramuscular *Hs6st2*/*Hs3st5* co-transduction in the TA of DMD rats. **a**, Representative tibialis anterior crosssections from wild-type (WT), vehicle-treated Del52, and co-transduced (CoT) Del52 rats, immunolabeled for GFP (reporter for the empty vector, or for *Hs3st5* under co-transduction) (in green), mCherry (*Hs6st2* reporter) (in red) , and DAPI (in blue); merged images below. Open arrowheads, reporter-positive myonuclei; arrows, centronucleated fibers. Scale bar, 1 mm, all panels. **b**, Percentage of transduced fibers. **c**, Representative Hs4c3 and DAPI labeling. Arrows, basal lamina; open arrowheads, vascular labeling; asterisks, cytoplasmic labeling. **d,** Global Hs4c3 labeling intensity. **e**, Percentage of centronucleated fibers. **f**, Representative laminin (LAMA1, in green) immunolabeling, color-coded cross-sectional area (CSA) intervals, and collagen VI immunolabeling (in red) as a marker of extracellular matrix deposition and fibrosis. g, Total fiber number. **h**, CSA distribution as percentage of total fibers per interval (inset, 8,000-12,000 μm² detail), with percentage of fibers below 500 μm² and fiber perimeter. **i**, Fibrotic area percentage. ns, not significant; *P < 0.05; **P < 0.01.

## Notes

### Competing Interest Statement

The authors have declared no competing interest.

